# Characterization of human lightness discrimination thresholds for independent spectral variations

**DOI:** 10.1101/2023.06.16.545355

**Authors:** Devin Reynolds, Vijay Singh

## Abstract

The lightness of an object is an intrinsic property that depends on its surface reflectance spectrum. The visual system estimates an object’s lightness from the light reflected off its surface. The light reflected also depends on object extrinsic properties of the scene. For stable perception, the visual system needs to discount variations due to extrinsic properties. We characterize this perceptual stability for variation in two spectral properties of the scene: the reflectance spectra of background objects and the intensity of light sources. We use a two-alternative forced-choice task to measure human observers’ thresholds of discriminating computer-generated images of 3D scenes based on the lightness of a spherical target object in the scene. We measured how the discrimination thresholds changed as we varied the reflectance spectra of the objects and the intensity of the light sources in the scene, both individually and simultaneously. For small amounts of extrinsic variations, the thresholds of discrimination remained constant indicating that the thresholds were dominated by observers’ intrinsic representation of lightness. As extrinsic variation increased, it started affecting observers’ lightness judgment and the thresholds increased. We estimated that the effects of extrinsic variations were comparable to observers’ intrinsic variation in the representation of object lightness. Moreover, for simultaneous variation of these spectral properties, the increase in threshold square compared to no variation condition was a linear sum of the corresponding increase in threshold squares for the individual properties, indicating that the variation from these independent sources combines linearly.

## INTRODUCTION

Our visual system provides perceptual representations of distal properties of objects based on the proximal stimuli captured by the eyes. While object properties are intrinsic to the object (its color, shape, etc.), the proximal stimuli also depend on the properties of the scene in which the object lies (such as background objects in the scene, illumination, etc.) as well as the position and pose of the observer. The task of the visual system is to provide stable correlates of object intrinsic properties under variability of the proximal signal due to object extrinsic scene properties. This work quantifies the extent to which the visual system provides such stability for the representation of the reflectance of an object under variation in spectral properties of the scene, specifically, variation in the spectra of the background objects and the intensity of light sources in the scene.

The perceptual correlate of the diffuse spectral reflectance of an object is its perceived color. For achromatic objects, the analogous perceptual quantity is object lightness. The human visual system is known to provide a relatively stable representation of the color/lightness of an object despite variability in the proximal signal due to changes in the light source, the surface reflectance of objects in the scene, and the geometry and other properties of the scene (Foster, 2011; Brainard & Radonjic, 2004). The degree to which such stability can be achieved is termed color/lightness constancy (Adelson, 2000; Gilchrist, 2006). Human color/lightness constancy has been measured using appearance-based approaches and discrimination-based approaches (Olkkonen & Ekroll, 2016). Appearance-based approaches involve tasks in which the observer makes a judgment about the appearance of stimuli. This approach includes methods such as color matching, color naming, scaling, and nulling (Foster, 2003). In color matching, observers adjust a test stimulus to match a standard stimulus. Color matching experiments show varying degrees of constancy with constancy measured between 15%-90% under conditions such as changes of illumination (Arend & Goldstein, 1987; Arend & Spehar, 1993), reflectance (Arend & Spehar, 1993; Patel, Munasinghe, & Murray, 2018), illumination gradients (Arend & Goldstein, 1990; Brainard, Brunt, & Speigle, 1997), and illumination and simulated reflectance (Rutherford & Brainard, 2002). Color naming is a more direct and arguably natural method to measure color constancy where observers are asked to categorize stimuli based on their hue, saturation, and lightness (Troost & De Weert, 1991). This method has been used with real (Uchikawa, Uchikawa, & Boynton, 1989; Olkkonen, Witzel, Hansen, & Gegenfurtner, 2010) and simulated stimuli (Olkkonen, Hansen, & Gegenfurtner, 2009) to measure constancy. Color naming methods have the limitation that there is a vast number of possible discernible colors (Linhares, Pinto, & Nascimento, 2008), but there is a limit to the gamut that can be displayed. Typically, observers are asked to name from a small set of colors (Speigle & Brainard, 1996; Smithson & Zaidi, 2004; Hansen, Walter, & Gegenfurtner, 2007) which might provide an overestimate of the measured constancy. In color scaling methods, observers view a stimulus and provide a rating on a scale for a set of colors, thus allowing for a finer level of comparison for measuring constancy (Luo, et al., 1991; Schultz, Doerschner, & Maloney, 2006). Scaling methods can also be used to measure changes in stimuli, where observers provide a rating of the change between stimuli (Ennis & Doerschner, 2019). Nulling or achromatic adjustment methods involve changing a test stimulus such that it appears achromatic (Arend, 1993; Brainard, 1998; Delahunt & Brainard, 2004). This method has the limitation that it provides data only for achromatic/gray stimuli and additional assumptions about the observers’ criterion need to be made for color appearances (Speigle & Brainard, 1996).

Discrimination-based approaches provide an objective method to measure color constancy (Bramwell & Hurlbert, 1996; Reeves, Amano, & Foster, 2008). In these experiments, observers discriminate stimuli as to whether they are the same or different from each other. The stimuli are varied in some relevant parameter space to measure the threshold for discriminating changes in the parameter (Craven & Foster, 1992; Pearce, Crichton, Mackiewicz, Finlayson, & Hurlbert, 2014; Aston, Radonjic, Brainard, & Hurlbert, 2019). Recently, Singh et. al (Singh, Burge, & Brainard, 2022) developed an equivalent noise paradigm that relates thresholds of discrimination to the variability in observers’ intrinsic representation of object properties (e.g., its lightness) and the variability due to object extrinsic properties of the scene.

They measured human lightness discrimination thresholds as a function of the amount of variability in the spectra of background objects in a scene. They related the discrimination thresholds to the variance in observers’ internal perceptual representation of lightness and the variance in the spectrally induced extrinsic variability. A comparison of the strength of intrinsic and extrinsic variability provided a measure of the degree of constancy in object intrinsic property due to the variability in object extrinsic property.

This equivalent noise paradigm can also be used to compare the effect of different sources of variabilities. The variance of multiple extrinsic properties can be characterized relative to the variance of intrinsic variability. These in turn can be compared to each other to measure their relative effects. It can also be used to characterize how the effect of multiple sources of variability combines when presented simultaneously.

In this work, we use this paradigm to compare the variation in two spectral properties of the scene to human observers’ representation of lightness. The spectral variations we study are the surface reflectance of the background objects in the scene and the intensity of the light sources in the scene. We measure human observers’ threshold of discriminating two images based on the lightness of an achromatic target object in the images. We measure how these discrimination thresholds change as we increase the variability in the reflectance spectra of the background objects and the intensity of the light sources. We measure discrimination thresholds for individual and simultaneous variations of these properties. We use the equivalent noise paradigm to relate the thresholds to the variance of observers’ intrinsic noise and extrinsic variability. These variances allow one to compare the relative effect of these spectral variations. A comparison of variances of individual and simultaneous variation conditions can provide information about the combination rules of multiple sources of variation.

We show that as the variability in the extrinsic sources increases, initially for a small amount of variation, the thresholds remain constant. In this region, the thresholds are determined primarily by the variation in observers’ internal representation of lightness. As the variability increases further, the discrimination thresholds increase. The increase in thresholds can be accounted for by a model based on signal detection theory. This model shows that the effect of extrinsic variation is within a factor of two compared to the variability in the intrinsic representation of lightness. This confirms that the visual system provides a large degree of lightness constancy under object extrinsic scene variation. By comparing the increase in thresholds of the individual and simultaneous variation condition from the no extrinsic variation condition, we show that the effects of individual sources combine linearly under simultaneous variation.

The paper is organized as follows. Section 2, Experimental Methods, provides the details of the experimental methods, stimuli used, and model fitting. Section 3, Results, provides the results of three experiments: variation in background reflectance spectra, variation in light source intensity, and simultaneous variation in these two properties. Section 4, Discussion, provides a summary of the results and remarks.

## 2 EXPERIMENTAL METHODS

### Overview

We followed the methodology published previously by (Singh, Burge, & Brainard, 2022). In this previous work, human lightness discrimination thresholds were measured under variability of the reflectance spectra of background objects in the scene. The work presented here follows the same experimental methods, except that the stimuli used in the experiments are different. This section provides an overview of the methods, focusing on the differences from the previous work. We refer the reader to the previous work for details.

Similar to the previous work, we used a two-alternative forced-choice (2AFC) procedure to measure thresholds (Figure 1). On each trial of the experiment, observers viewed pairs of computer-generated 3D scenes displayed on a color-calibrated monitor. These pairs consisted of a standard image and a comparison image. The images were presented sequentially for 250ms, with a 250ms inter-stimulus interval between them. Both images contained a centrally located achromatic sphere as the target object. Observers were required to indicate the image on which the target object was lighter. Between trials, we manipulated the luminous reflectance factor (LRF) of the target object in the comparison image. The LRF is defined as the ratio of the luminance of a surface under a reference illuminant and the luminance of the reference illuminant itself (American Society for Testing and Materials, 2017). The reference illuminant was chosen as the CIE D65 standard illuminant. The order of the standard and the comparison image was chosen in a pseudorandom order. We plotted the psychometric functions of the observers by collecting data on the proportion of times the observer chose the comparison image to be lighter (see Figure 2 for an example). To estimate observers’ discrimination threshold, we fit the proportion comparison chosen data with a cumulative normal function. We defined the threshold as the difference between the LRF of the target object for which the cumulative normal fit was equal to 0.76 and 0.50. This corresponds to a d-prime of 1 in a two-interval task.

**Figure 1:**
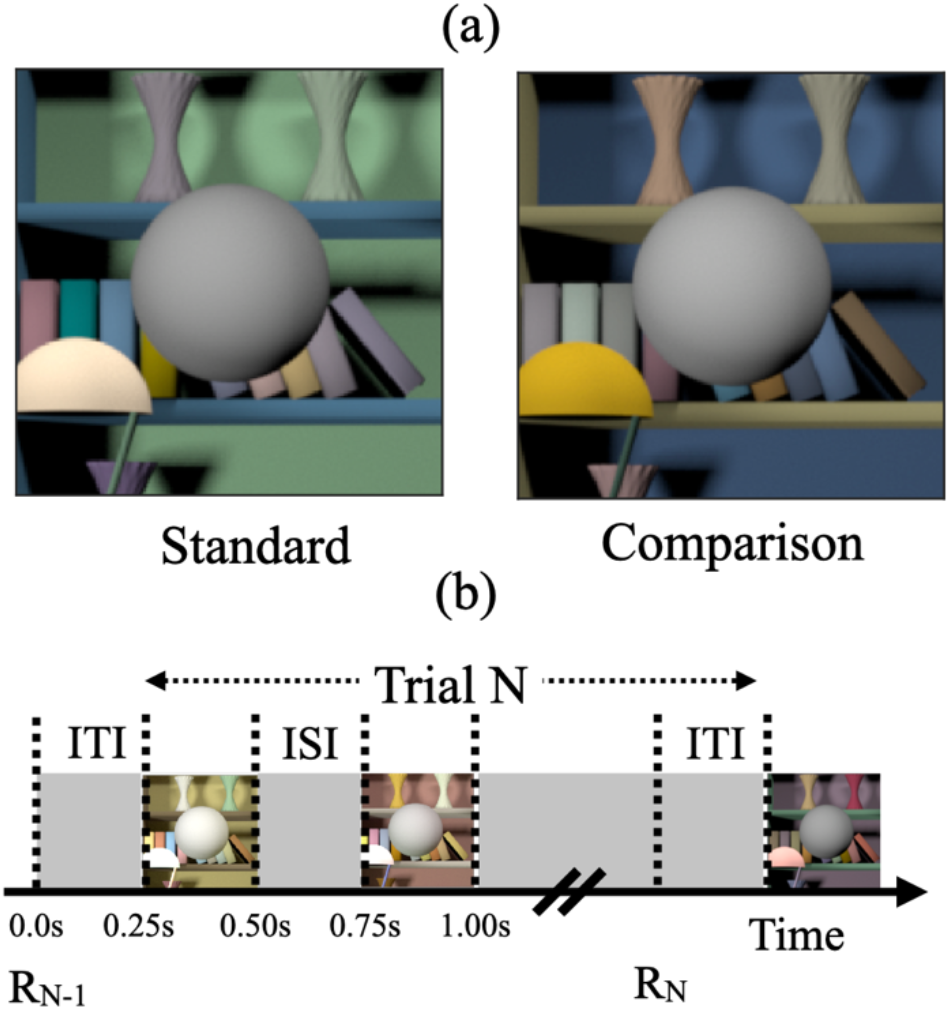
(a) Psychophysical task. (Adapted from Figure 1 in Singh, Burge, & Brainard, 2022) The psychophysical task involved comparing two images, a standard image and a comparison image, on each trial and selecting the image with the lighter target object. The target object was an achromatic sphere at the center of the image. The images were generated computationally by graphically rendering models of 3D scenes. They were displayed on a color-calibrated monitor. This panel shows examples of standard and comparison images. The reflectance spectrum of the target object was spectrally flat, and the target object appeared gray. The reflectance of the target object in the standard image was held fixed and it changed for the comparison image. In this panel, the target object in the comparison image is lighter. We measured the fraction of times the observers chose the target object in the comparison image to be lighter as a function of the lightness of the target object in the comparison image. The fraction comparison chosen data was used to determine the lightness discrimination threshold (Figure 2). We studied how the lightness discrimination thresholds changed as the trial-to-trial variability in the reflectance spectra of the background objects and the intensity of the light sources increased. **(b) Trial sequence:** R_N-1_ indicates the recording of the observer’s response for the (N-1)^th^ trial. The N^th^ trial begins 250ms after the completion of the (N-1)^th^ trial (Inter Trial Interval, ITI = 250ms). In the N^th^ trial, the standard and comparison images are presented for 250ms each with a 250ms inter-stimulus interval (ISI) in between the two images. The order of the standard and comparison images is chosen in pseudorandom order. The observer records their choice by pressing a button on a gamepad after both images have been presented and removed from the screen. The observers could take as long as they wish before making their choice. The recording of their choice is indicated by R_N_ in the panel. The next trial begins 250ms after the choice has been recorded.

**Figure 2:**
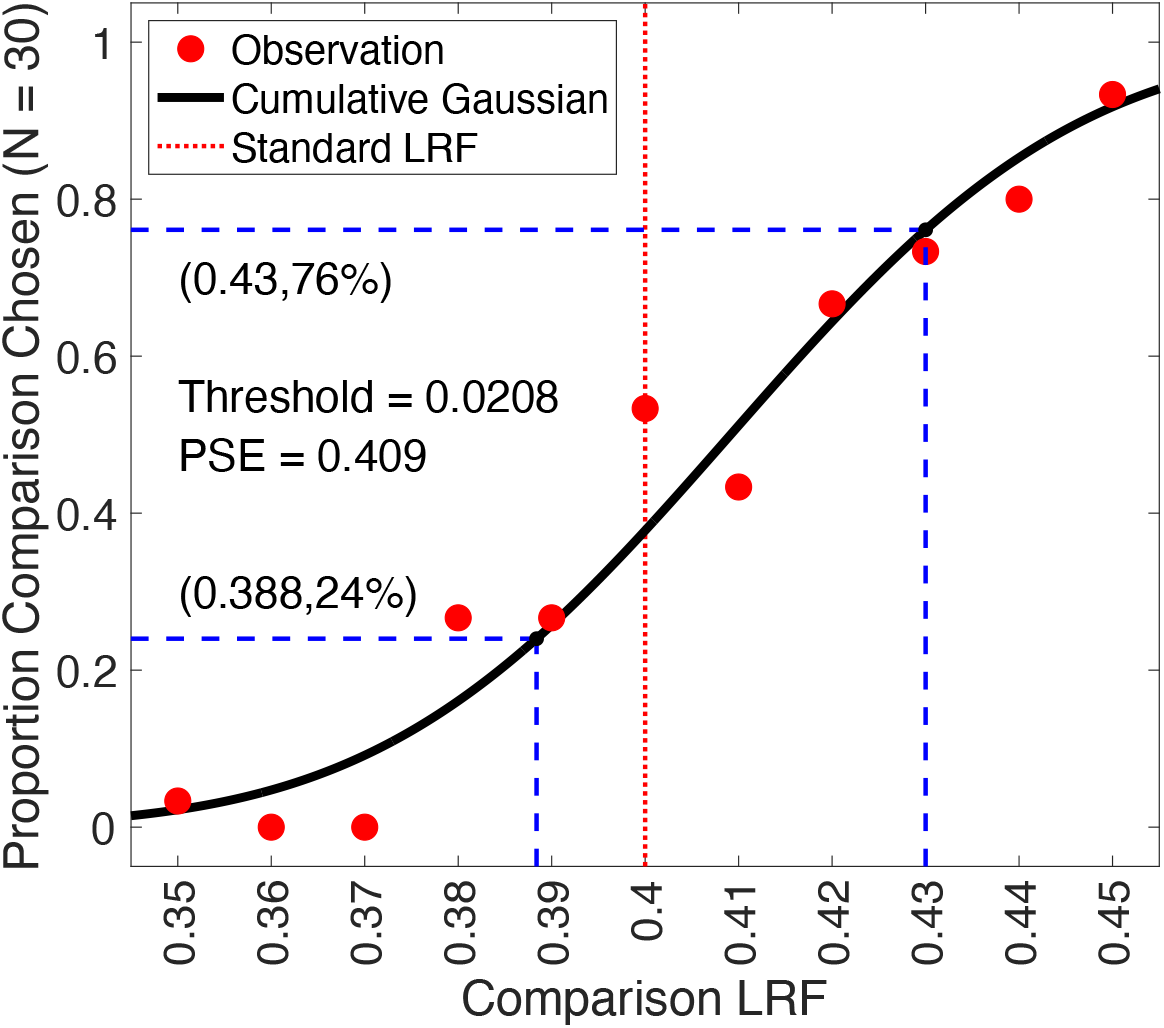
Psychometric function: We measured the proportion of times the observers selected the target in the comparison image to be lighter as a function of the LRF (lightness reflectance factor) of the target object. We collected 30 responses for each of the 11 equally spaced values of the comparison image target object LRF, ranging from 0.35 to 0.45. The LRF of the target object in the standard image was 0.40. The LRF of the target object in the comparison image was selected in a pseudorandom order. To analyze the data, we used maximum likelihood methods to fit a cumulative normal function to the proportion comparison chosen data. We imposed constraints on the guess rate and lapse rate, requiring them to be equal and within the range of 0 to 0.05. The threshold was determined as the difference between the LRF values corresponding to a proportion comparison chosen of 0.76 and 0.50, obtained from the cumulative normal fit. The figure presented here illustrates the data for observer 0003 in the second block of the background reflectance variation experiment (previously registered as Experiment 6) for the no variation (σ^2^ = 0.00, δ = 0.00) condition. The discrimination threshold was measured to be 0.0208. The point of subjective equality (PSE), which corresponds to a proportion of 0.5 in the comparison task, was found to be 0.409. The lapse rate for this particular fit was determined to be 0.00.

We measured the effect of variation in two types of object-extrinsic scene properties on human lightness discrimination thresholds: variation in the reflectance spectra of the background objects in the scene and variation in the intensity of the light sources in the scene. We performed three experiments. These experiments were preregistered (see below Preregistration).

(1) <u>Background reflectance spectra variation (preregistered as Experiment 6):</u> In this experiment, we measured human lightness discrimination thresholds as a function of the amount of variation in the background objects while the spectra of the light sources were kept fixed.
(2) <u>Light source intensity variation (preregistered as Experiment 7):</u> In this experiment, we measured lightness discrimination thresholds as a function of the amount of variation in the intensity of the light sources while the background was fixed.
(3) <u>Simultaneous variation (preregistered as Experiment 8):</u> In this experiment, we measured lightness discrimination thresholds as both the background object reflectance spectra and the light source intensity varied simultaneously.

To study the effect of variation in surface reflectance of background objects we generated samples of surface reflectance spectra from a multivariate normal distribution. The distribution was a statistical model of databases of natural surface reflectance measurements. The amount of variation in the surface reflectance of background objects was varied by changing the size of the covariance matrix of the multivariate normal distribution. We measured discrimination thresholds for both chromatic and achromatic variations. In the chromatic variation, the reflectance spectra could take any shape and the objects varied in their luminance and chromaticity. In the achromatic variation, the reflectance spectra were spectrally flat, and the objects were gray.

The shape of the spectral power distribution function of the light source was chosen as the CIE D65 reference illuminant. The intensity was varied by multiplying the spectral power distribution function by a scalar sampled from a log uniform distribution. The amount of variation was controlled by changing the range of the log uniform distribution.

The subsections below provide additional methodological detail.

### Preregistration

We preregistered the experiments performed in this work before collecting the data. The preregistration documents are available at: https://osf.io/7tgy8/.^1^ The documentation includes information about the experimental design as well as the procedure to estimate thresholds from the collected data.

The experiments were preregistered as Experiment 6 (referred to as Background reflectance spectra variation), Experiment 7 (referred to as Light source intensity variation), and Experiment 8 (referred to as Simultaneous variation). Experiment 6 was a replication of previous work (preregistered as Experiment 3; Singh, Burge, & Brainard, 2022) with additional conditions in which the background objects were achromatic and varied only in their lightness. While the stimuli were different for the three experiments (preregistered Experiments 6, 7, and 8), the experimental method to measure lightness discrimination thresholds was the same.

The preregistration documents mentioned that the experiments aimed at characterizing the dependence of human lightness discrimination thresholds on the amount of variation in the background reflectance and the intensity of the light source in the scene. The method of estimating discrimination thresholds was described in the document. We predicted that the thresholds would increase with an increase in the amount of variation. For background variation, we predicted that the thresholds of achromatic variation would be lower than chromatic variation. We also predicted that the increase in thresholds could be captured by an equivalent noise model (Singh, Burge, & Brainard, 2022). Additionally, we predicted that the threshold for simultaneous variation would be higher than the threshold for individual variations.

### Reflectance and Illumination Spectra

We used a statistical model of natural reflectance datasets to generate reflectance spectra of background objects (Singh, Cottaris, Heasly, Brainard, & Burge, 2018; Singh, Burge, & Brainard, 2022). We combined two datasets of surface reflectance measurements (Vrhel, Gershon, & Iwan, 1994; Kelly, Gibson, & Nickerson, 1943). These datasets contain 632 surface reflectance measurements. We mean-centered the dataset by subtracting out the mean surface reflectance over the 632 measurements. Then we used principal component analysis (PCA) to obtain the projection of the mean-centered dataset along the eigenvectors associated with the six largest eigenvalues. These eigenvalues captured more than 99.5% of the variance (Singh, Cottaris, Heasly, Brainard, & Burge, 2018). The empirical distribution of the projection weights thus obtained was approximated with a multivariate normal distribution. To get the projection weights of random samples of reflectance spectra, pseudorandom samples were generated from this multivariate normal distribution. Reflectance spectra were constructed by using these projection weights along with the eigenvectors and adding the mean of the surface reflectance dataset. A physical realizability condition was imposed on these spectra by ensuring that the reflectance at each wavelength was between 0 and 1. If a reflectance spectrum did not meet this criterion, it was discarded.

To generate achromatic surface reflectance spectra, after generating a physically realizable reflectance spectrum, its average reflectance over all wavelengths was calculated and it was replaced by a spectrum that had this average reflectance at all wavelengths.

To control the amount of variation in the reflectance spectra, the covariance matrix of the multivariate normal distribution was multiplied by a covariance scalar (σ^2^). A covariance scalar of 0 indicates that there is no variation in the reflectance spectra of the background objects. On the other hand, a covariance scalar of 1 corresponds to the range of reflectance variation observed in the combined natural reflectance datasets.

The light source power spectrum was chosen to be the CIE D65 reference illuminant. The D65 spectrum was divided by its mean power over wavelength to obtain its relative spectral shape. The variation in the light source intensity was introduced by multiplying the normalized D65 spectrum by a random sample generated from a log-uniform distribution in the range [1− δ, 1+ δ], where δ determines the range of the distribution. We chose log-uniform distribution for the multiplication parameter because the spectral power distribution function of natural daylight spectra varies over three orders of magnitude and their mean over wavelength can be roughly approximated by a log-uniform distribution (Singh, Cottaris, Heasly, Brainard, & Burge, 2018). All light sources in a scene were assigned the same power spectrum.

The values of the two parameters σ^2^ and δ for the three experiments were as follows:

<u>Background object reflectance variation (Experiment 6):</u> In this experiment, we generated images for nine conditions. Six of these conditions were for chromatic variation at six logarithmically spaced values of the covariance scalar (σ^2^): [0, 0.01, 0.03, 0.1, 0.3, 1.0] (same as Singh, Burge, & Brainard, 2022). Three conditions were for achromatic variation at covariance scalar (σ^2^): [0.03, 0.3, 1.0]. The power spectrum of the light source was the same for all images. The power spectrum multiplication scalar was assigned an arbitrary value of 5. Figure 3 shows five typical images for the nine conditions.

**Figure 3:**
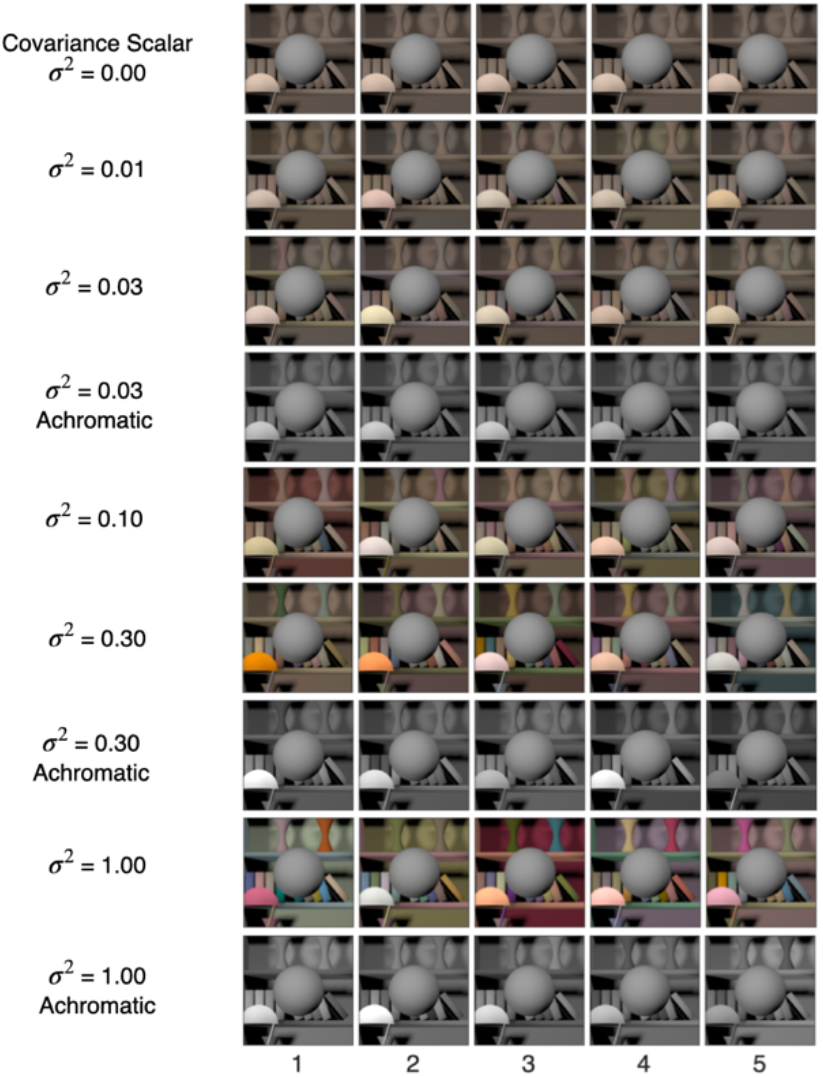
Background object reflectance variation: We studied two types of variations in the reflectance spectra of background objects in the scene: chromatic variation and achromatic variation. In the chromatic variation, the reflectance spectra could take any shape, and the objects varied in their luminance and chromaticity. In the achromatic variation, the reflectance spectra were spectrally flat, and the objects appeared gray and varied only in their luminance. The spectra were chosen from a multivariate normal distribution that modeled the statistics of natural reflectance spectra. The covariance matrix of the multivariate normal distribution was multiplied by a scalar to control the variance in the samples. We generated images at six logarithmically spaced values of the covariance scalar for chromatic variation and at three values of the covariance scalar for achromatic variations. The figure shows five typical images for each of these nine conditions. For each condition, we generated 1100 images, 100 images at 11 linearly spaced values of target object LRF in the range [0.35, 0.45]. The target object in each image in the figure is at LRF = 0.4.

<u>Light source intensity variation (Experiment 7):</u> In this experiment, we generated images for seven linearly spaced values of the range parameter (δ): [0.00, 0.05, 0.10, 0.15, 0.20, 0.25, 0.30]. The reflectance spectra of all background objects were the same and were equal to the mean spectrum of the reflectance database. This corresponds to a covariance scalar of 0. Figure 4 shows five typical images for the seven conditions.

**Figure 4:**
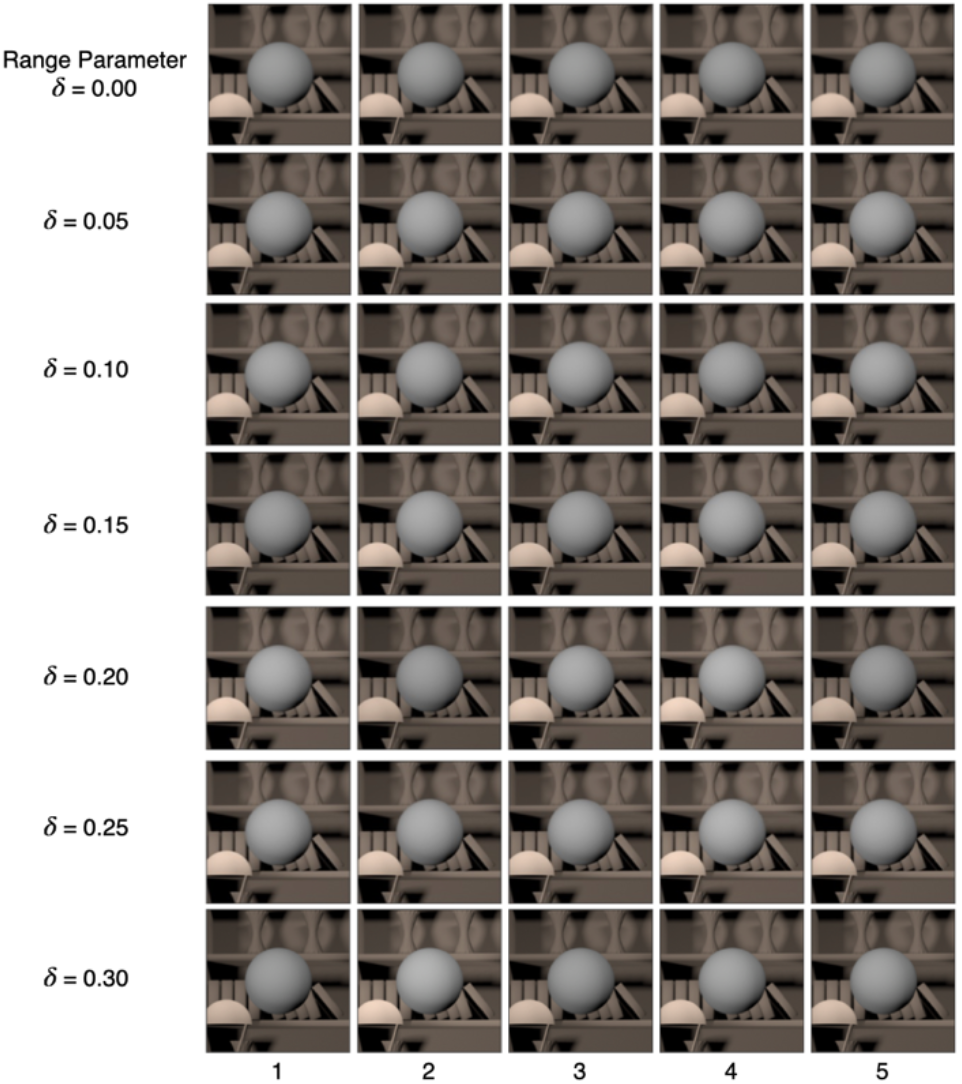
Light intensity variation: The shape of the power spectrum of the light sources in the scene was chosen to be CIE reference illuminant D65. The intensity of the power spectrum was varied by multiplying the normalized D65 spectrum with a scalar sampled from a log uniform distribution in the range [1-δ, 1+ δ]. The amount of variation was controlled by changing the value of the range parameter δ. We generated images at seven linearly spaced values of the range parameter in the range [0.00, 0.30]. For each value of the range parameter, we generated 1100 images, 100 images at each value of the target object LRF in the range [0.35, 0.45]. The figure shows five sample images at each of the seven values of the range parameter. The target object in each image in the figure has the same LRF of 0.40.

<u>Simultaneous variation (Experiment 8):</u> In this experiment, we studied six conditions. These were: no variation (σ^2^ = 0, δ = 0), chromatic background variation (covariance scalar = 1, δ = 0), achromatic background variation (σ^2^ = 1, δ = 0), light source intensity variation (σ^2^ = 0, δ = 0.3), simultaneous variation chromatic background (σ^2^ = 1, δ = 0.3) and simultaneous variation achromatic background (σ^2^ = 1, δ = 0.3). Figure 5 shows five typical images for these six conditions.

**Figure 5:**
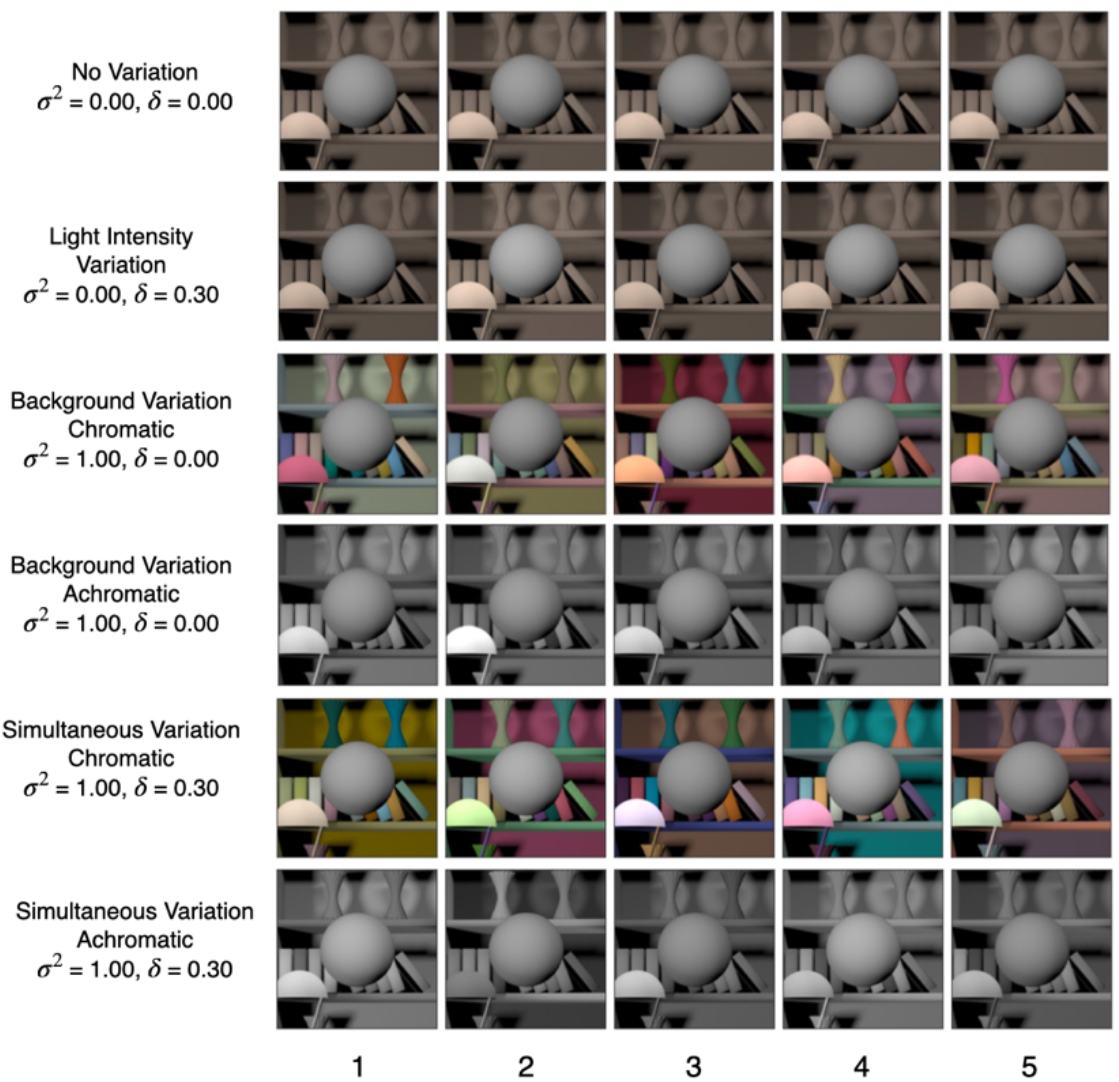
Simultaneous variation: This figure shows five sample images for the six conditions studied in preregistered Experiment 8. We generated 1100 images for each of these conditions, 100 images at each value of the target object LRF in the range [0.35, 0.45].

### Stimulus Design

The images used in this work were generated using the software Virtual World Color Constancy (VWCC) (github.com/BrainardLab/VirtualWorldColorConstancy) described in (Singh, Cottaris, Heasly, Brainard, & Burge, 2018). The initial step to generate an image involves constructing a 3D model that serves as the base scene. Then, based on the specific experimental condition, we assign reflectance spectra and spectral power distribution functions to the objects and light sources within the base scene. All light sources within a given scene were assigned identical spectral power distribution functions. Subsequently, we utilize Mitsuba, a physically-realistic open-source rendering system (mitsuba-renderer.org; Jakob, 2010) to produce a 2D multispectral image of the scene. A 201-pixel by 201-pixel part of the image centered at the target object was cropped out to display on the monitor. Monitor calibration data was used to convert the multispectral images to gamma-corrected RGB images as described in (Singh, Burge, & Brainard, 2022). Gamma-corrected RGB images were presented on the calibrated monitor during the experiment. When displayed on the experimental monitor at a distance of 75cm from the observers’ head position, the image was nearly 2° in visual angle with the target nearly 1° in visual angle.

For each condition described above, we generated 1100 images, 100 images at each of the 11 linearly spaced values of the target object LRF in the range [0.35, 0.45]. The standard image target object LRF was 0.4. The comparison image target object LRF varied in the range [0.35, 0.45]. We generated 100 images at each comparison level to avoid excessive replication of images in the experiment. For the no variation (σ^2^ = 0.00, δ = 0.00) condition, we generated one image at each target object LRF level, as the background reflectance and the light source intensity remained fixed in this case. The scene geometry remained fixed during the experiment and the images did not include secondary reflections.

The standard image, when presented on the experimental monitor, had an average luminance of 87.1 cd/m^2^ for the condition (σ^2^ = 0.00, δ = 0.00). The average luminance of the target object for the 11 LRF levels were [120.9, 122.3, 123.8, 125.2, 126.5, 127.9, 129.2, 130.5, 131.9, 133.1, 134.4] cd/m^2^.

For the (σ^2^ = 1.00, δ = 0.30) condition, the average luminance of the standard image was 87.8 cd/m^2^. The average luminance of the target object for the 11 LRF levels were [117.7, 119.4, 119.4, 122.3, 123.7, 123.8, 127.8, 126.9, 127.7, 129.1, 129.0] cd/m^2^.

### Experimental Structure

In this study, a trial is defined as the display of a standard and a comparison image on the monitor and the recording of the observer’s response. An interval is defined as the presentation of either the standard image or the comparison image within a trial. A block consists of recording 330 trials for one condition, 30 trials at each of the 11 comparison image target LRF levels. A permutation consists of recording one block of data for each condition in an experiment. We recorded three permutations for each observer in each experiment. Each permutation had a random order of the blocks.

The order of the blocks in a permutation, the LRF levels of the comparison image in trials of a block, and the order of standard and comparison images in a trial were generated pseudorandomly and stored at the beginning of the experiment for each observer. Before starting a new permutation for an observer, the data for all blocks (conditions) in a permutation was collected.

A session consisted of recording three blocks on a single day. An observer performed no more than one session a day. Each block in a session was divided into three sub-blocks of 110 trials. Between these sub-blocks, the observers took a break of a minimum one-minute duration. The observers also took a small break (nearly two to five minutes) between blocks. The observers were informed that they could stop the experiment at any time. If the observer terminated a block, the data was not recorded. No observer terminated a block of the experiment.

Each observer first performed a practice session where three blocks of data were recorded for the no variation (σ^2^ = 0, δ = 0) condition. The observers were excluded from the experiment if their mean threshold for the last two blocks was higher than 0.03. If the observer passed this criterion, then the rest of the data was collected over several days.

The experimental procedure was explained to each observer at the beginning of the practice session. The experimenter then obtained the consent for the experiment. Vision tests were performed on the observer to ensure normal visual acuity and normal color vision. After this, the observer went to the experimental room where they were familiarized with the experimental set-up by performing a familiarization block of 40 trials. Then the observer was dark adapted by sitting in the dark for about 5 minutes. Then the data for the three blocks of the practice session was recorded. At the end of the practice session, the observer was informed if they could continue the experiment.

If the observer was continued, their data was collected over several sessions. The data for all six observers of an experiment was collected over several weeks. The data of all six observers for preregistered Experiment 6 was collected before starting preregistered Experiment 7. The data of all six observers for preregistered Experiment 7 was collected before starting preregistered Experiment 8.

### Observer Recruitment and Exclusion

The study recruited observers from North Carolina Agricultural and Technical State University and the local Greensboro community. The participants were compensated for their time. The observers underwent a screening process to meet criteria such as a normal visual acuity of 20/40 or better (with corrective eyewear if necessary) and normal color vision, which was assessed using pseudo-isochromatic plates (Ishihara, 1977). The preregistration documents outlined these exclusion criteria (see Methods: Preregistration).

We further conducted a practice session to identify observers who could reliably perform the psychophysical task. In the practice session, the observer performed three blocks of the experiment for the no variation condition (σ^2^ = 0.00, δ = 0.00) and the threshold was calculated for these three blocks. If the mean threshold of the observer for the last two blocks in the practice session was larger than 0.030 (log T^2^, –3.2), the observer was discontinued. The preregistration document specified this exclusion criterion (See Methods: Preregistration).

If the observer met these criteria, they continued with the rest of the experiment.

For each observer, the practice session was performed at the beginning of each of the three experiments (Experiments 6, 7, and 8), irrespective of whether the observer had participated in an earlier experiment.

### Observer Information

<u>Background reflectance variation (preregistered Experiment 6):</u> A total of 25 observers participated in the practice sessions for the background reflectance variation experiment (10 Females, 15 Males; age 19-34; mean age 22.9). Observers were given pseudo-names to deidentify their personal information from the data. Six of these observers (pseudo-names: *0003, bagel, committee, content, observer*, and *revival*) met the performance criterion set for screening (2 Females, 4 Males; age 19-28; mean age 23.33). Their visual acuity assessed using the Snellen chart was 20/40 or better in both eyes and color vision assessed using Ishihara plates was normal. The visual acuities of the observers in the main experiment were: *0003*, L = 20/30, R = 20/20; *bagel*, L = 20/20, R = 20/20; *committee*, L = 20/25, R = 20/25; *content*, L = 20/20, R = 20/20; *observer*, L = 20/25, R = 20/25; *revival*, L = 20/20, R = 20/20. *Committee, content,* and *observer* wore personal corrective eyewear both during vision testing and during the experiments. Observers *0003, bagel*, and *revival* did not require or use corrective eyewear.

<u>Light source intensity variation (preregistered Experiment 7):</u> A total of 15 observers participated in the practice sessions for the light source intensity variation experiment (9 Females, 6 Males; age 19-33; mean age 25). Six of these observers (pseudo-names: *0003*, *bagel*, *content*, *oven*, *primary*, and *revival*) met the performance criterion set for screening (3 Females, 3 Males; age 19-28; mean age 23.83). Their visual acuity assessed using the Snellen chart was 20/40 or better in both eyes and color vision assessed using Ishihara plates was normal. The visual acuities of the observers in the main experiment were: *0003*, L = 20/30, R = 20/30; *bagel*, L = 20/20, R = 20/20; *content*, L = 20/20, R = 20/20; *oven*, L = 20/20, R = 20/20; *primary*, L = 20/20, R = 20/20; *revival*, L = 20/20, R = 20/20. Observer *content* and *primary* wore personal corrective eyewear both during vision testing and during the experiments. Observers *0003, bagel*, *oven,* and *revival* did not require or use corrective eyewear. Observer *oven* reported some difficulties during a few sessions of the experiment and their thresholds for two conditions did not fit the expected pattern. We removed their data from the analysis presented in this work. Their data and thresholds are provided as supplementary information (SI).

<u>Simultaneous variation (preregistered Experiment 8):</u> A total of 20 observers participated in the practice sessions for the simultaneous variation experiment (9 Females, 11 Males; age 19-28; mean age 20.8). Six of these observers (pseudo-names: *0003, bagel, content, oven, manos,* and *revival*) were retained for the experiment (2 Females, 4 Males; age 19-28; mean age 23.33). Four observers (*0003, bagel, content,* and *oven*) met the screening criteria specified in the preregistration. Due to a lack of observers who met the preregistration criteria, two observers (*manos,* and *revival*), whose thresholds were close to the preregistration criteria, were also retained for the experiment. Observer *Revival* had participated in the previous two experiments and had met the criteria both times. Observer *Manos* showed improvement in thresholds with each block, with the threshold for the final block below 0.03. This was a deviation from the preregistration. Their visual acuity assessed using the Snellen chart was 20/40 or better in both eyes and color vision assessed using Ishihara plates was normal. The visual acuities of the observers in the main experiment were: *0003*, L = 20/30, R = 20/30; *bagel*, L = 20/20, R = 20/20; *content*, L = 20/20, R = 20/20; *oven*, L = 20/20, R = 20/20; *manos*, L = 20/25, R = 20/25; *revival*, L = 20/20, R = 20/20. Observer *Content* wore personal corrective eyewear both during vision testing and during the experiments. Observers *0003, bagel*, *manos*, *oven*, and *revival* did not require or use corrective eyewear.

### Apparatus

The experiments were performed in a dark room and the stimuli were presented on a color-calibrated LCD monitor (27-in. NEC MultiSync EA271U; NEC Display Solutions). The pixel resolution of the monitor was selected as 1920 x 1080. Its refresh rate was 60Hz and each RGB channel was operated at 8-bit resolution. The experimental computer was an Apple Macintosh with an Intel Core i7 processor. The experimental programs were written in MATLAB (MathWorks; Natick, MA) and utilized Psychophysics Toolbox (http://psychtoolbox.org) and mgl (http://justingardner.net/doku.php/mgl/overview) libraries. A Logitech F310 gamepad controller was used to collect observers’ responses.

The distance between the observers’ eyes and the monitor was set at 75cm. A forehead rest and chin cup (Headspot, UHCOTech, Houston, TX) were used to stabilize the observers’ head position. The observers’ eyes were centered both horizontally and vertically in relation to the display.

### Monitor Calibration

The monitor was calibrated using a spectroradiometer (PhotoResearch PR655) as described in (Singh, Burge, & Brainard, 2022). The monitor was calibrated before starting each experiment. Once calibrated, the same settings were used till data for all observers for that experiment was collected. The monitor was then recalibrated for the next experiment. Data was collected in the sequence Experiment 6, Experiment 7, and Experiment 8.

### Ethics Statement

All experiments were approved by North Carolina Agricultural and Technical State University Institutional Review Board and were in accordance with the World Medical Association Declaration of Helsinki.

### Code and Data Availability

The data for each experiment and observer is provided as supplementary information (SI). The SI is available at: https://github.com/vijaysoophie/SimultaneousVariationPaper. The SI contains the proportion comparison chosen data as well as the thresholds for the 3 experimental blocks of each condition, for each experiment and observer. The MATLAB scripts to generate Figures 2, 6 – 12, supplementary Figures S1-S7, and the scripts to obtain thresholds of the linear receptive field formulation of the model are also provided in the SI.

### Linear Receptive Field Model

The thresholds of preregistered Experiment 6 were fit to the linear receptive field model developed by (Singh, Burge, & Brainard, 2022). This model consisted of a simple center-surround receptive field (*R*). The receptive field was square in shape to match the images in the psychophysics experiment. Its center was a circle of radius equal to the size and the location of the target object. The central region had a spatially uniform positive sensitivity of 1. The surround had a spatially uniform negative sensitivity of v_s_. The receptive field response was computed as the dot product of the receptive field with the standard and the comparison images. A mean zero Gaussian noise was added to the response. The image with the higher noise-added receptive field response was chosen to be lighter. The two parameters of the model, noise variance (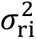) and surround sensitivity (v_s_), provided an estimate of the internal noise (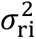) and the variance of the extrinsic properties (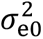). The model related the thresholds (*T*) in the experiments to the variance in the intrinsic noise (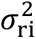) of the observer and the extrinsic variance (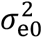) through the relation:

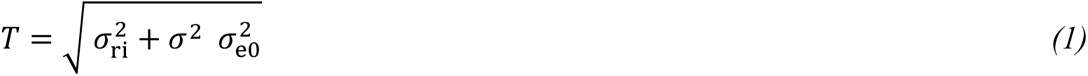

where σ^2^ is the covariance scalar (see (Singh, Burge, & Brainard, 2022) for details). The variance of the extrinsic properties (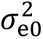) resulting from the variation in the image can be computed as 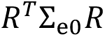, where 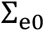 is the covariance matrix of the variation in the images. We calculate the pixel-by-pixel covariance matrix of the image database for each condition and use the receptive field vector *R* to estimate the quantity 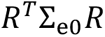.

<u>Background reflectance variation:</u> To estimate the variance of the intrinsic noise of the observer and extrinsic variation in the images, we chose the value of the Gaussian noise variance (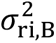) and the surround sensitivity v_s,B_ to minimize the mean squared difference between the model and experimental thresholds measured at the six values of the covariance scalar. 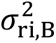 provided the estimate of the intrinsic noise. The extrinsic noise (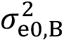) was estimated by using the best-fit surround sensitivity (v_s,B_) of the receptive field (*R*) and the sample covariance matrix of the images (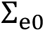) at σ^2^ = 1 in the relation 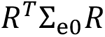.

<u>Light source intensity variation:</u> We fit a functional form similar to Eq. (1) to the thresholds of the light source intensity variation experiment (Experiment 7) where we replaced the covariance scalar σ^2^ by the range parameter δ. We chose the value of the Gaussian noise variance (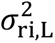) and receptive field surround sensitivity (v_s,L_) to minimize the mean square difference between the observer and model thresholds measured at the seven values of the range parameter. 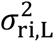 provided the estimate of the observer’s intrinsic noise.

In natural viewing conditions, the intensity of the light source varies over several orders of magnitude (Singh, Cottaris, Heasly, Brainard, & Burge, 2018). This range of variation cannot be captured in the experiment due to the limitations of the monitor. To estimate the extrinsic noise, we used the best-fit surround sensitivity (v_s,L_) and the sample covariance matrices to calculate the quantity 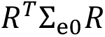 as a function of the range parameter δ. We fit the resulting values with an exponential function (see Figure S1). The extrinsic noise (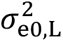) was chosen as the value of the exponential fit at δ = 1. The parameter (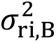) could be used to estimate the extrinsic noise for more naturalistic variations using the exponential fit.

## 3 RESULTS

### Human Lightness Discrimination Thresholds Increase with Background Reflectance Variation

We measured the lightness discrimination thresholds of human observers for two types of variation in the reflectance spectra of background objects in the scene: chromatic variation and achromatic variation. In the chromatic variation, the reflectance spectra could take any shape and thus the background objects varied in their chromaticity and luminance. In the achromatic variation, each spectrum had the same reflectance at all wavelengths, and thus the spectra varied only in their overall luminance and the objects were gray. The reflectance spectra were sampled from a statistical model of natural surface reflectance spectra databases (See Methods: Reflectance and Illumination Spectra). The model approximated the database with a multivariate normal distribution whose covariance matrix was multiplied by a covariance scalar (σ^2^) to control the amount of variation in the sampled spectra. We measured the discrimination thresholds of six human observers at six values of the covariance scalar for chromatic variation and three values of covariance scalar for achromatic variation. For each of the nine conditions and each observer, discrimination thresholds were measured three times in three separate blocks. The psychometric functions for these nine conditions are shown for one observer in Figure 6. The psychometric functions of all six observers are shown in Figure S3. We notice that as the covariance scalar increases, the slope of the psychometric functions decreases, corresponding to an increase in discrimination thresholds. The thresholds for chromatic and achromatic variation are comparable.

**Figure 6:**
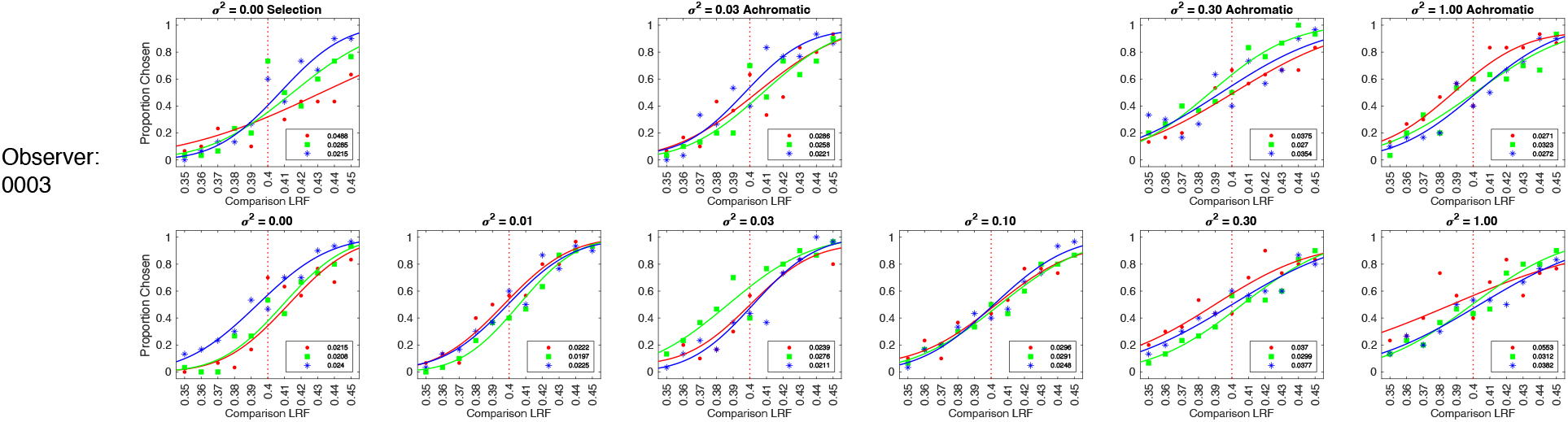
Psychometric functions for observer 0003 for background reflectance variation experiment: We measured the proportion comparison chosen data for the nine conditions separately in three blocks for each observer. The figure shows the psychometric function for observer 0003. The psychometric functions for all six observers are shown in Figure S2. A cumulative normal function was fit to the data from each block to determine the discrimination threshold (see Figure 2). The legend provides the estimated lightness discrimination threshold for each block, obtained from the cumulative fit. The first panel in the top row shows the data and thresholds for the selection session. The selection session was a practice session in which the threshold for the no variation condition was measured three times. An observer was selected for the experiment only if the average of their last two discrimination threshold measurements in the selection session was less than 0.30. The last three panels in the top row show the data for the three achromatic conditions. The bottom row shows the data for the chromatic variation conditions.

Figure 7 shows the change in discrimination thresholds as a function of the variance in the reflectance of the background objects. Here we plot the mean log threshold squared (averaged across observers, N = 6) as a function of the log of the covariance scalar. The thresholds and standard error of the mean (SEM) from Figure 7 are listed in Table S1. We observe that the thresholds are nearly constant at small values of the covariance scalar. Once the value of the covariance scalar starts increasing, the log threshold squared increases. Further, the thresholds are comparable for chromatic and achromatic variation. The p-values of the hypothesis that the mean thresholds for chromatic and achromatic variations are equal are 0.72, 0.57, and 0.16 for covariance scalar 0.03, 0.30, and 1.00 respectively, indicating that the differences in the mean thresholds are not statistically significant. Figure S3 shows the discrimination thresholds measured in preregistered Experiment 6 and previously reported thresholds by (Singh, Burge, & Brainard, 2022). The measured thresholds are consistent between the two experiments.

**Figure 7:**
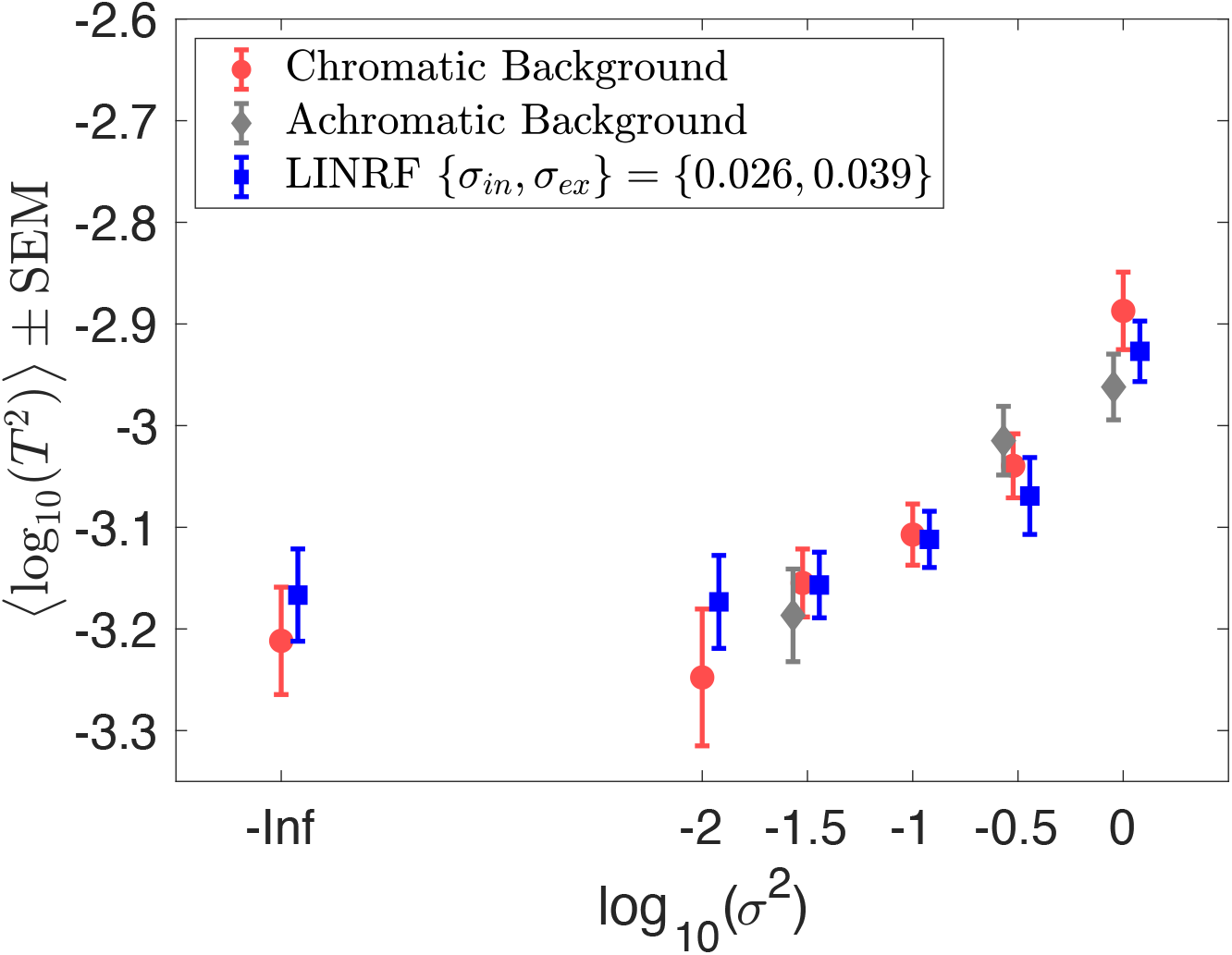
Background variation increases lightness discrimination thresholds. Mean (N = 6) log squared threshold vs log covariance scalar from human psychophysics for chromatic (red circles) and achromatic conditions (gray diamonds). The error bars represent +/-1 SEM taken between observers. The threshold of the linear receptive field (LINRF) model was estimated by simulation for the six values of the covariance scalar (blue squares). The blue error bars show +/-1 standard deviation estimated over 10 independent simulations. The legend shows the parameters of the linear receptive field (LINRF) model fit. The data has been jittered for ease of viewing. A comparison of the thresholds with the previously published data in Singh, Burge, Brainard 2022 is shown in Figure S3.

We fit the thresholds to the linear receptive field (LINRF) model (Eq. 1) developed by (Singh, Burge, & Brainard, 2022). The LINRF model provides the estimate of the variance of the internal noise of the observer as 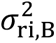 = 0.026 and the variance of the extrinsic variability due to the reflectance of background objects as 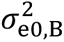 = 0.039. The equivalent noise level, the ratio of the external variance to intrinsic noise, is ∼ 1.5, indicating that the variability in the representation of object lightness induced by the natural variability in the reflectance of background objects is close to the internal variability of that representation. If the ratio was equal to 1, then we would have concluded that the visual system has discounted the external variability. But the ratio is not significantly large compared to 1, this indicates that the visual system provides a significant level of lightness constancy.

### Human Lightness Discrimination Thresholds Increase with Light Source Intensity Variation

We measured the lightness discrimination thresholds of human observers as we varied the intensity of light sources in the scene. The shape of the spectrum of the light sources was fixed to be standard daylight spectrum D65. We normalized the spectrum by its mean over wavelengths. The intensity was varied by multiplying the normalized spectrum by a scalar sampled from a log-uniform distribution in the range [1-δ, 1+ δ]. The reflectance spectra of the background objects were fixed. We measured lightness discrimination thresholds for seven values of the range parameter δ for five human observers. The psychometric functions of one of the observers for these seven conditions are shown in Figure 8. The psychometric functions of all six observers are shown in Figure S4. Figure 9 shows how the thresholds change as the amount of variation in the light source intensity increases. The data is averaged over five observers (also see Figure S5). Table S2 lists the mean thresholds and the SEM measured in this experiment. Similar to the trend for reflectance spectra variation, lightness discrimination thresholds remain constant for small values of the range parameter, and then the log threshold squared increases with an increase in the range parameter. A fit of the mean squared threshold with the linear receptive field model gives the value of internal noise as 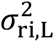 = 0.028. This compares well with the internal noise obtained from the background reflectance spectra variation experiment (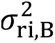 = 0.026). The variance of the extrinsic variability estimated at the range parameter δ = 1.00 is 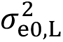 = 0.052. The equivalent noise level, the ratio of external variation to intrinsic noise is ∼ 1.8. This indicates that the variation in the lightness representation induced by the variation in light source intensity is close to the internal variation of that representation at these levels. In natural conditions, where light source intensity varies over several orders of magnitude, the extrinsic noise can be estimated by generating images at the level of variation and using the LINRF model.

**Figure 8:**
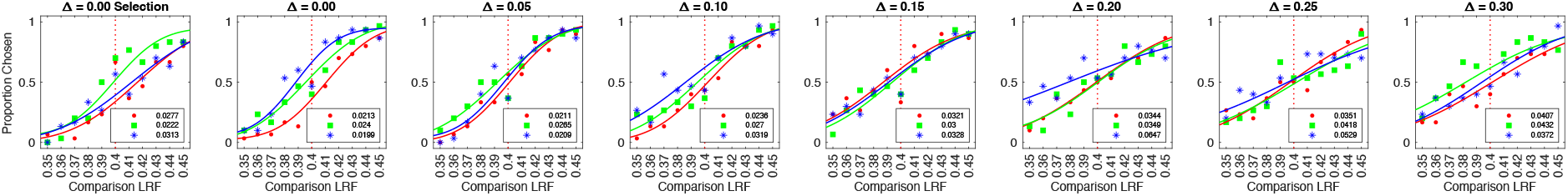
Psychometric functions for observer 0003 for light intensity variation experiment: Same as Figure 6, but for the light intensity variation experiment. The figure shows the proportion comparison chosen data for the selection session and the seven conditions for observer 0003. The psychometric functions for all observers are shown in Figure S4.

**Figure 9:**
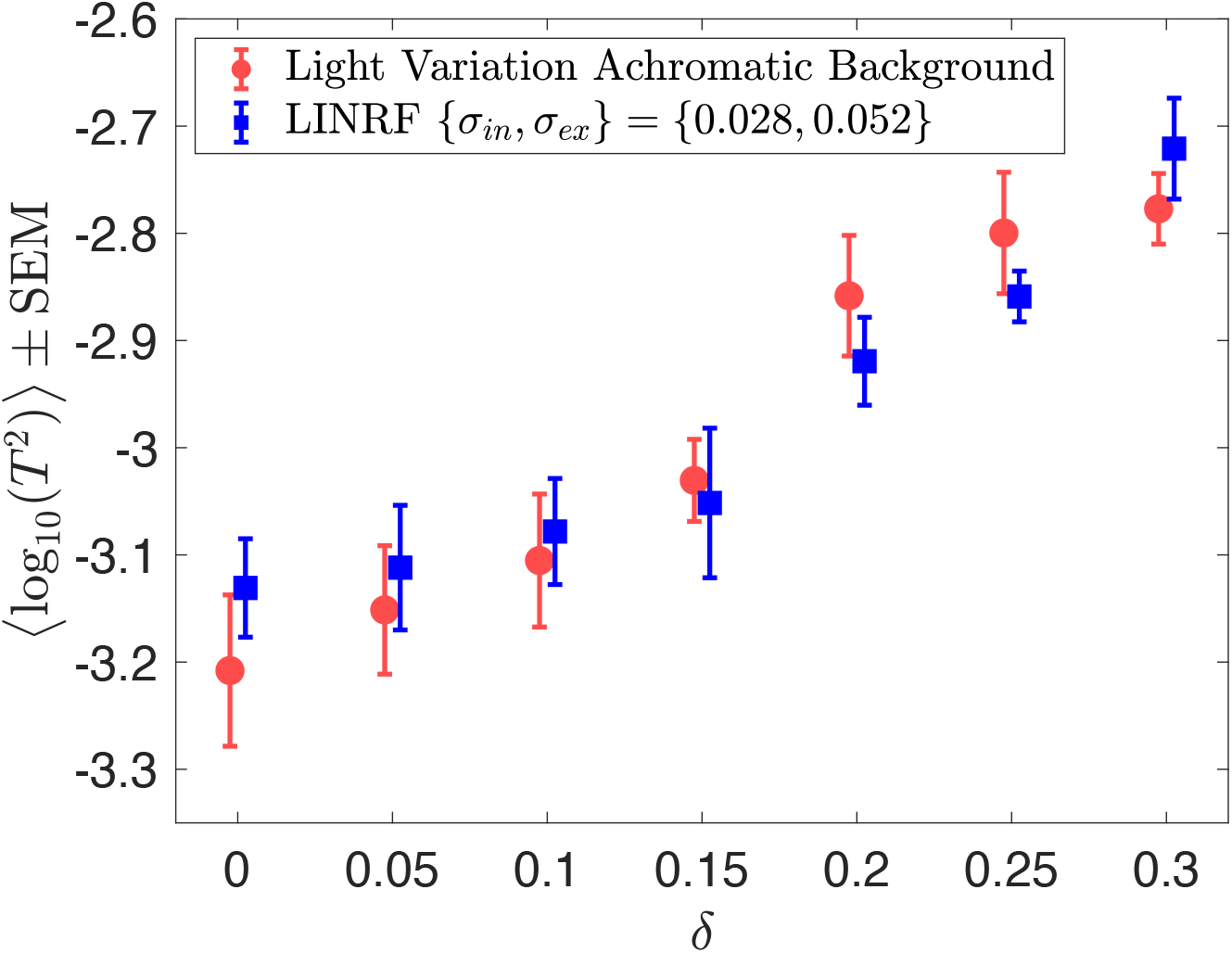
Light source intensity variation increases lightness discrimination threshold. Mean (N = 5) log squared threshold vs range parameter from human psychophysics for the seven light source intensity variation conditions (red circles). The error bars represent +/-1 SEM taken between observers. The threshold of the linear receptive field (LINRF) model was estimated by simulation for the seven values of the range parameters (blue squares). The blue error bars show +/-1 standard deviation estimated over 10 independent simulations. The legend shows the parameters of the LINRF model fit. The data has been jittered for ease of viewing. The data for all six observers is shown in Figure S5.

### Thresholds for Simultaneous Variation are Higher Than Individual Variations

We measured lightness discrimination thresholds for simultaneous variation in the reflectance spectra of background objects and the intensity of the light sources in the scene. In this experiment, we studied six conditions: no variation, variation in the reflectance spectra of background objects with a fixed spectrum of the light sources for achromatic and chromatic backgrounds, variation in the intensity of light sources with a fixed background, and simultaneous variation in the intensity of the light sources and the reflectance spectra of background objects for chromatic and achromatic backgrounds. We measured the lightness discrimination thresholds of six human observers for these six conditions. The psychometric function of one of the observers is shown in Figure 10. The psychometric functions of all six observers are shown in Figure S6. Figure 11 shows the mean log squared threshold of all six observers for these six conditions. Table S3 lists the mean thresholds and SEM from Figure 11. The threshold for simultaneous variation of light source intensity and reflectance spectra of background objects is higher than the condition with individual variations in these properties. As observed earlier, the threshold for achromatic and chromatic conditions are comparable. The p-value of the hypothesis that the mean thresholds for chromatic and achromatic variations are equal is 0.19 for the background variation condition and 0.44 for the simultaneous variation condition, indicating that the differences in the mean thresholds are not statistically significant.

**Figure 10:**
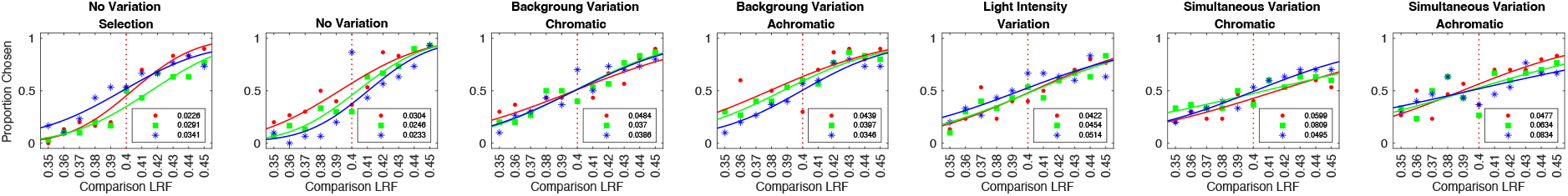
Psychometric functions for observer 0003 for simultaneous variation experiment: Same as Figures 6 and 8, but for simultaneous variation experiment. The figure shows the proportion comparison chosen data for the selection session and the six conditions for observer 0003. The data for all observers are shown in Figure S6.

**Figure 11:**
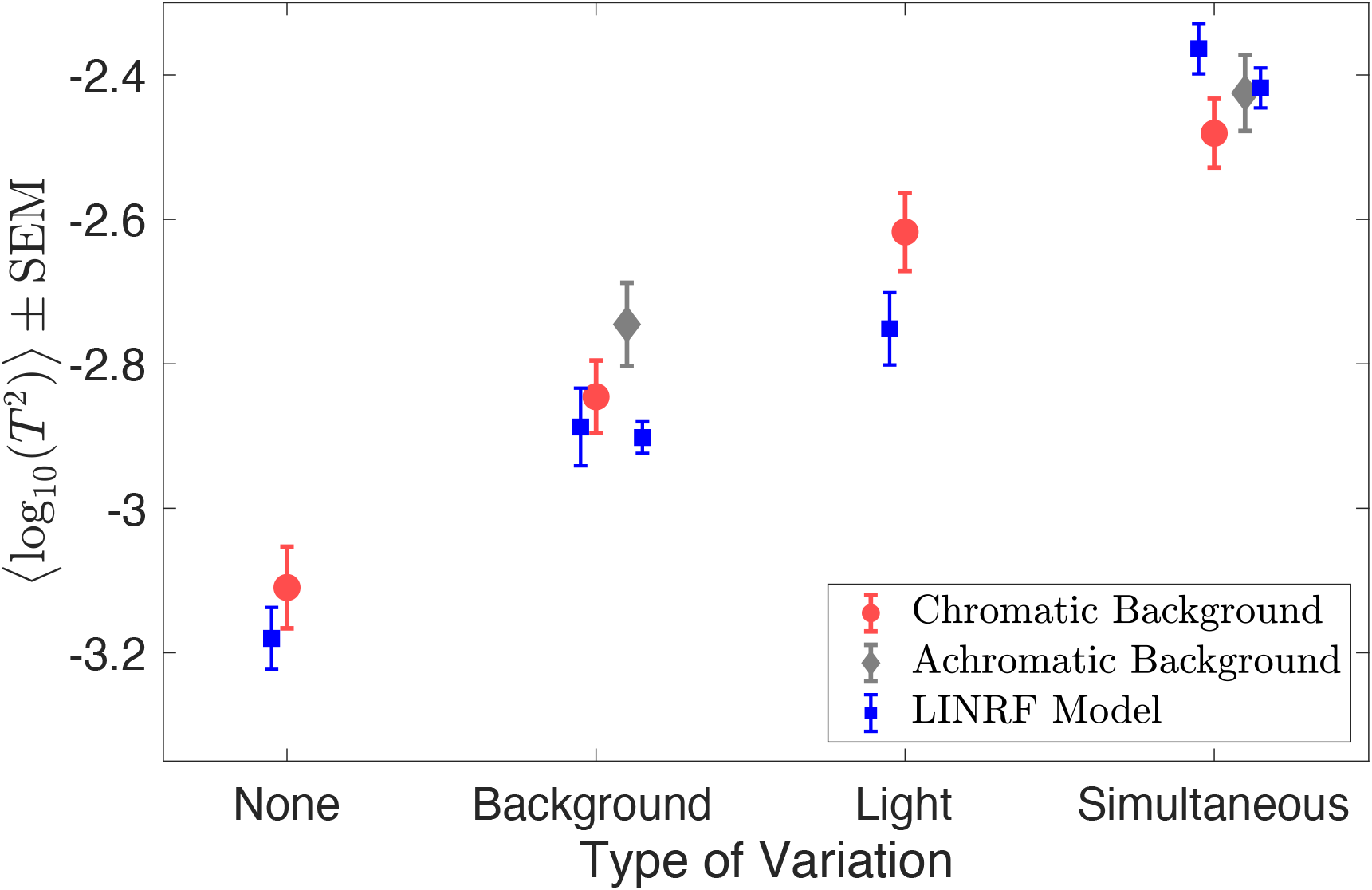
Discrimination thresholds for simultaneous variation of two sources are higher than individual discrimination thresholds. Mean (N = 6) log squared threshold for the six conditions in simultaneous variation experiment. The error bars represent +/-1 SEM taken between observers. The data for chromatic (red circles) and achromatic (gray diamonds) conditions have been plotted next to each other for visual comparison. The thresholds of the linear receptive field (LINRF) model (blue squares) were estimated using the parameters of the background variation condition (Figure 7) for the None, Background variation, and Simultaneous variation conditions and using the parameters of the light intensity variation condition (Figure 9) for the Light condition. The blue error bars show +/-1 standard deviation estimated over 10 independent simulations. See Figure S7 for LINRF model thresholds with the same set of parameters for all conditions.

Figure 11 also shows the squared thresholds of the LINRF model for the six conditions. We used the intrinsic noise and the surround sensitivity (v_s_) parameters of the background reflectance variation experiment to estimate the threshold of the LINRF model for the no-variation condition, background spectra variation conditions, and simultaneous variation conditions (Experiment 6, Figure 7). For the light intensity variation condition, we used the parameters of the light source intensity variation experiment (Experiment 7, Figure 9). See Figure S7 for the predictions of the LINRF model with the same set of parameters for all six conditions.

We can use the linear receptive field model to compare the extrinsic variance of the simultaneous variation condition to the variance of the individual variations. According to the linear receptive model, the square of the threshold is proportional to the sum of the variance of observers’ intrinsic noise and the extrinsic variation in the stimuli (Eq. 1). The squared threshold at the no variation condition is equal to the variance of the observers’ intrinsic noise. In the case of extrinsic variation, the increase in threshold square compared to the no variation condition equals the variance of the extrinsic variation. When there is more than one independent source of extrinsic variation, the total variance of the simultaneous variation should be the sum of the variance of the individual variations. This predicts that the increase in threshold square for simultaneous variation conditions should be equal to the sum of the corresponding increase for the individual variation conditions.

Figure 12 shows the increase in the mean squared threshold above the no variation condition. We compare the mean square thresholds of the simultaneous variation condition with the sum of the mean square thresholds of the individual conditions for chromatic and achromatic conditions. The increase in the squared threshold of the simultaneous variation condition is comparable to the sum of the increase in the squared threshold for the individual variations. The p-value of the hypothesis that the mean increase in squared thresholds for simultaneous variation is equal to the sum of the mean increase in the squared thresholds of light intensity variation and background object reflectance variation are 0.86 and 0.80 for chromatic and achromatic conditions respectively. The variance of the extrinsic noise calculated for the background variation condition (σ^2^ = 1.00, δ = 0.00) is 0.0015 and the light intensity variation condition (σ^2^ = 0.00, δ = 0.30) is 0.0017. As expected, the variance of the simultaneous variation condition (σ^2^ = 1.00, δ = 0.30), which is 0.0033, is comparable to the sum of individual variances (0.0032).

**Figure 12:**
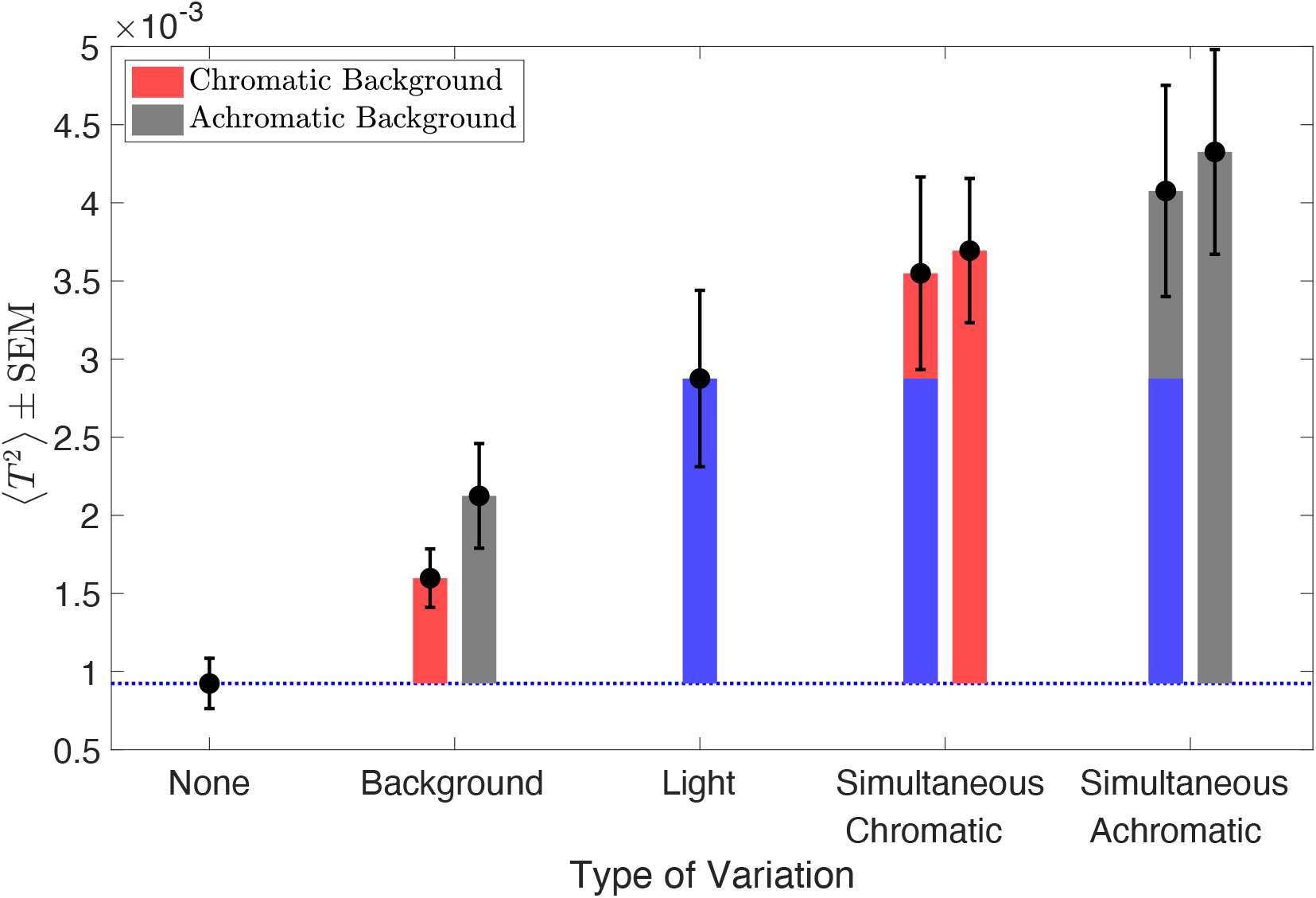
Extrinsic noise of independent variations add linearly for simultaneous variation: Mean squared thresholds (N=6) for the six conditions in simultaneous variation experiment (black circles). The black error bars represent +/-1 SEM taken between observers. The bars (red, gray, blue) represent the increase in squared thresholds compared to the no variation condition (blue dotted line). For the simultaneous variation conditions, the bars on the right (bars with one color, red or gray) represent the increase in the measured squared threshold and the bars on the left (stacked bars of two different colors) represent the increase in the sum of the squared threshold of the light intensity variation (blue bar) and the corresponding background variation conditions (red or gray).

## 4 DISCUSSION

The visual system is known to maintain a stable representation of object lightness despite variations in the proximal signal due to the scene. In this work, we characterize the extent of such stability for variability in the spectra of background objects and light sources in a scene. We measured human observers’ threshold of discriminating two objects based on their lightness as a function of the amount of variation in these spectral properties. We observe that for low levels of variability, the thresholds first remained constant, showing that in this regime the performance was determined by observers’ intrinsic noise. As the variability increased, the effect of extrinsic variation started dominating the performance and the discrimination thresholds increased. Using a model based on signal detection theory, we related the thresholds in the low variability regime to the internal noise of the observer. The model also related the increase in threshold to the amount of variability in the extrinsic property, thus providing a comparison of the variance in the extrinsic property to the intrinsic noise. The effects of both types of extrinsic variation, the spectra of background objects and the intensity of light sources, were comparable to the effect of intrinsic noise, showing that the visual system provides a good degree of constancy to these variations. Further, for simultaneous variation of these properties, the effects add linearly, resulting in the variance of the simultaneous variation to be equal to the sum of the variance of the individual variations.

**Chromatic v/s Achromatic Variations:** Lightness discrimination thresholds of chromatic and achromatic variation in the reflectance spectra of background objects were statistically similar. The chromatic aspect of the variation does not seem to influence lightness discrimination, indicating that lightness and chromaticity are encoded independently. This hypothesis could be tested by measuring chromaticity discrimination thresholds under chromatic and achromatic variation of background objects.

**Visual system at threshold level:** For the spectral variabilities studied in this work, the variances in observers’ representation of lightness due to extrinsic variations are within a factor of two compared to the variances in observers’ intrinsic representation of lightness. If these variances were equal, one could conclude that the visual system has fully compensated for the extrinsic variation. As the extrinsic variances are larger than the variance of the intrinsic noise, the visual system has not fully compensated for the external variabilities. But since these variances are within a factor or two, it shows that the visual system provides a large degree of stability in the perceptual representation of lightness and seems to work at near-threshold levels.

**Rules of Combination:** The increase in the squared threshold of simultaneous variation of reflectance spectra of background object and intensity of light sources from no variation condition was equal to the sum of the increase in the squared threshold of the individual variations. This could be accounted for by assuming that the sources of noise are independent, and their effects add linearly.

## 5 ACKNOWLEDGEMENTS

We thank Dr. David Brainard for his comments on the manuscript. This work was supported by the National Science Foundation award BCS 2054900.

**Table S1:** Thresholds for Background Variation Experiment (Preregistered Experiment 6): Mean threshold (averaged over blocks) ± SEM of six human observers for nine background variation conditions studied in preregistered experiment 6.

| Condition | Observer |  |  |  |  |  |
| --- | --- | --- | --- | --- | --- | --- |
|  | 0003 | Bagel | Committee | Content | Observer | Revival |
| $\sigma^2 = 0.00$ | 0.0221±0.0010 | 0.0185±0.0018 | 0.0344±0.0027 | 0.0223± 0.0012 | 0.0311±0.0053 | 0.0251±0.0023 |
| $\sigma^2 = 0.01$ | 0.0215±0.0009 | 0.0194±0.0020 | 0.0386±0.0103 | 0.0193±0.0012 | 0.0263±0.0059 | 0.0262±0.0048 |
| $\sigma^2 = 0.03$ | 0.0242±0.0019 | 0.0261±0.0020 | 0.0285±0.0029 | 0.0246±0.0046 | 0.0292±0.0007 | 0.0282±0.0016 |
| $\sigma^2 = 0.03$<br>Achromatic | 0.0255±0.0019 | 0.0213±0.0024 | 0.0343±0.0055 | 0.0227±0.0023 | 0.0267±0.0040 | 0.0263±0.0016 |
| $\sigma^2 = 0.10$ | 0.0278±0.0015 | 0.0238±0.0010 | 0.0284±0.0017 | 0.0278±0.0035 | 0.0335±0.0024 | 0.0281±0.0013 |
| $\sigma^2 = 0.30$ | 0.0348±0.0025 | 0.0277±0.0024 | 0.0344±0.0020 | 0.0286±0.0002 | 0.0277±0.0019 | 0.0301±0.0038 |
| $\sigma^2 = 0.30$<br>Achromatic | 0.0333±0.0032 | 0.0284±0.0028 | 0.0319±0.0047 | 0.0308±0.0015 | 0.0358±0.0030 | 0.0287±0.0022 |
| $\sigma^2 = 1.00$ | 0.0416±0.0072 | 0.0316±0.0008 | 0.0379±0.0024 | 0.0323±0.0022 | 0.0405±0.0042 | 0.0360±0.0055 |
| $\sigma^2 = 1.00$<br>Achromatic | 0.0289±0.0017 | 0.0310±0.0015 | 0.0391±0.0029 | 0.0384±0.0058 | 0.0312±0.0015 | 0.0322±0.0009 |

**Table S2.** Thresholds for Lightness Intensity Variation Experiment (Preregistered Experiment 7): Mean threshold (averaged over blocks) ± SEM of six human observers measured for seven lightness intensity conditions studied in preregistered experiment 7. The thresholds of observer Oven were not used in Figure 9.

| Condition | Observer |  |  |  |  |  |
| --- | --- | --- | --- | --- | --- | --- |
|  | 0003 | Bagel | Content | Oven | Primary | Revival |
| $\delta = 0.00$ | 0.0217±0.0012 | 0.0181±0.0001 | 0.0208±0.0014 | 0.0520±0.0114 | 0.0329±0.0061 | 0.0372±0.0008 |
| $\delta = 0.05$ | 0.0228±0.0018 | 0.0229±0.0018 | 0.0207±0.0007 | 0.0580±0.0064 | 0.0346±0.0042 | 0.0364±0.0013 |
| $\delta = 0.10$ | 0.0275±0.0024 | 0.0217±0.0009 | 0.0242±0.0040 | 0.0325±0.0022 | 0.0343±0.0013 | 0.0376±0.0072 |
| $\delta = 0.15$ | 0.0316±0.0009 | 0.0238±0.0011 | 0.0323±0.0032 | 0.0333±0.0019 | 0.0345±0.0042 | 0.0326±0.0002 |
| $\delta = 0.20$ | 0.0447±0.0100 | 0.0381±0.0046 | 0.0276±0.0016 | 0.0493±0.0120 | 0.0423±0.0050 | 0.0392±0.0034 |
| $\delta = 0.25$ | 0.0433±0.0052 | 0.0393±0.0062 | 0.0308±0.0023 | 0.0461±0.0060 | 0.0532±0.0083 | 0.0387±0.0025 |
| $\delta = 0.30$ | 0.0404±0.0018 | 0.0429±0.0033 | 0.0347±0.0014 | 0.0580±0.0061 | 0.0465±0.0047 | 0.0421±0.0042 |

**Table S3.** Thresholds for Simultaneous Variation Experiment (Preregistered Experiment 8): Mean threshold (averaged over blocks) ± SEM of six human observers measured for six conditions studied in preregistered experiment 8.

| Condition | Observer |  |  |  |  |  |
| --- | --- | --- | --- | --- | --- | --- |
|  | 0003 | Bagel | Content | Oven | Manos | Revival |
| No Variation | 0.0261±0.0022 | 0.0227±0.0019 | 0.0246±0.0004 | 0.0383±0.0066 | 0.0258±0.0036 | 0.0366±0.0085 |
| Background Variation Chromatic | 0.0414±0.0036 | 0.0340±0.0058 | 0.0392±0.0083 | 0.0498±0.0050 | 0.0306±0.0013 | 0.0383±0.0033 |
| Background Variation Achromatic | 0.0394±0.0027 | 0.0319±0.0015 | 0.0427±0.0074 | 0.0683±0.0048 | 0.0435±0.0071 | 0.0389±0.0010 |
| Light Intensity Variation | 0.0464±0.0027 | 0.0656±0.0208 | 0.0412±0.0021 | 0.0592±0.0091 | 0.0464±0.0046 | 0.0474±0.0069 |
| Simultaneous Variation Chromatic | 0.0635±0.0092 | 0.0536±0.0014 | 0.0437±0.0011 | 0.0639±0.0106 | 0.0768±0.0085 | 0.0528±0.0037 |
| Simultaneous Variation Achromatic | 0.0648±0.0103 | 0.0540±0.0017 | 0.0478±0.0049 | 0.0826±0.0166 | 0.0749±0.0082 | 0.0561±0.0028 |

---

**Figure S1:**
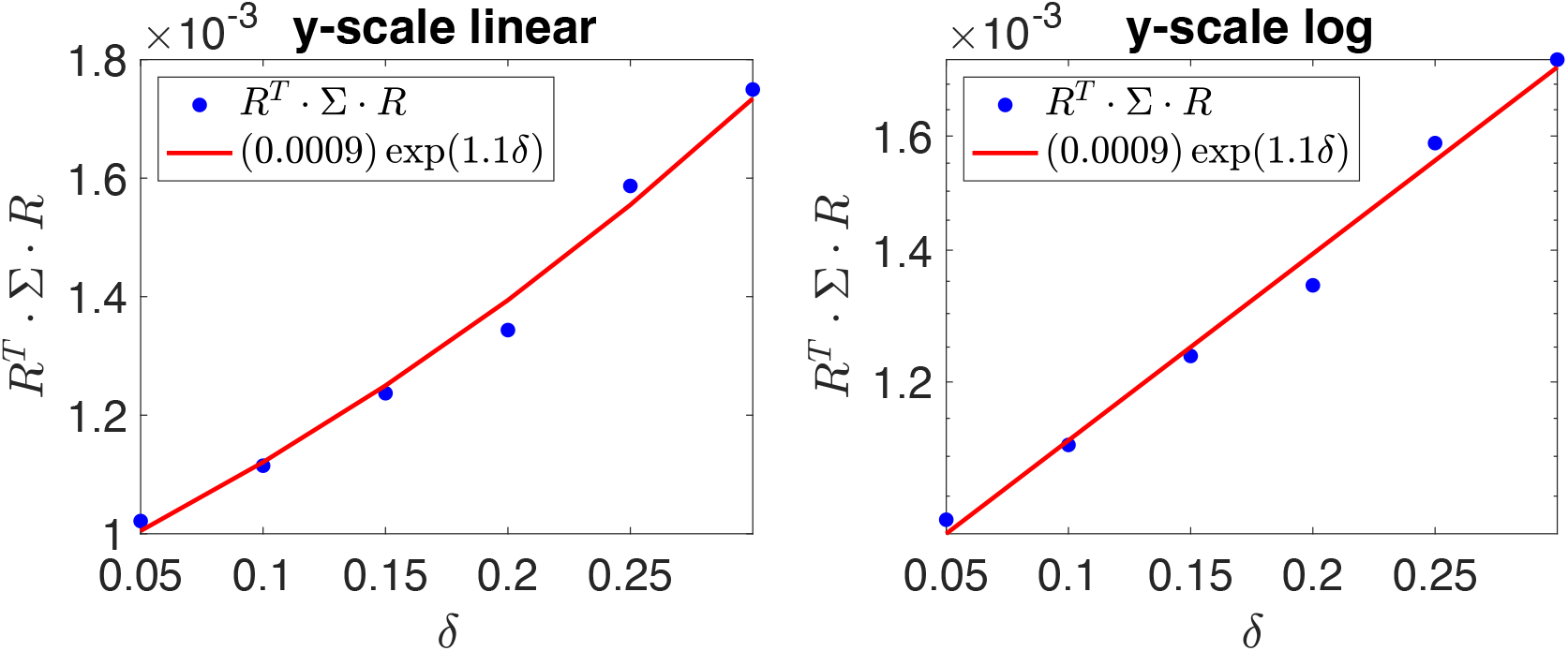
Estimation of extrinsic noise for light intensity variation: Plot of the variance (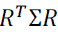) as a function of the range parameter δ on a linear (left panel) and logarithmic (right panel) scale. We fit the function with an exponential of the form *A* * exp (*B* ⋅ δ). The variance in the extrinsic noise is estimated as the value of the fit at δ = 1.

**Figure S2:**
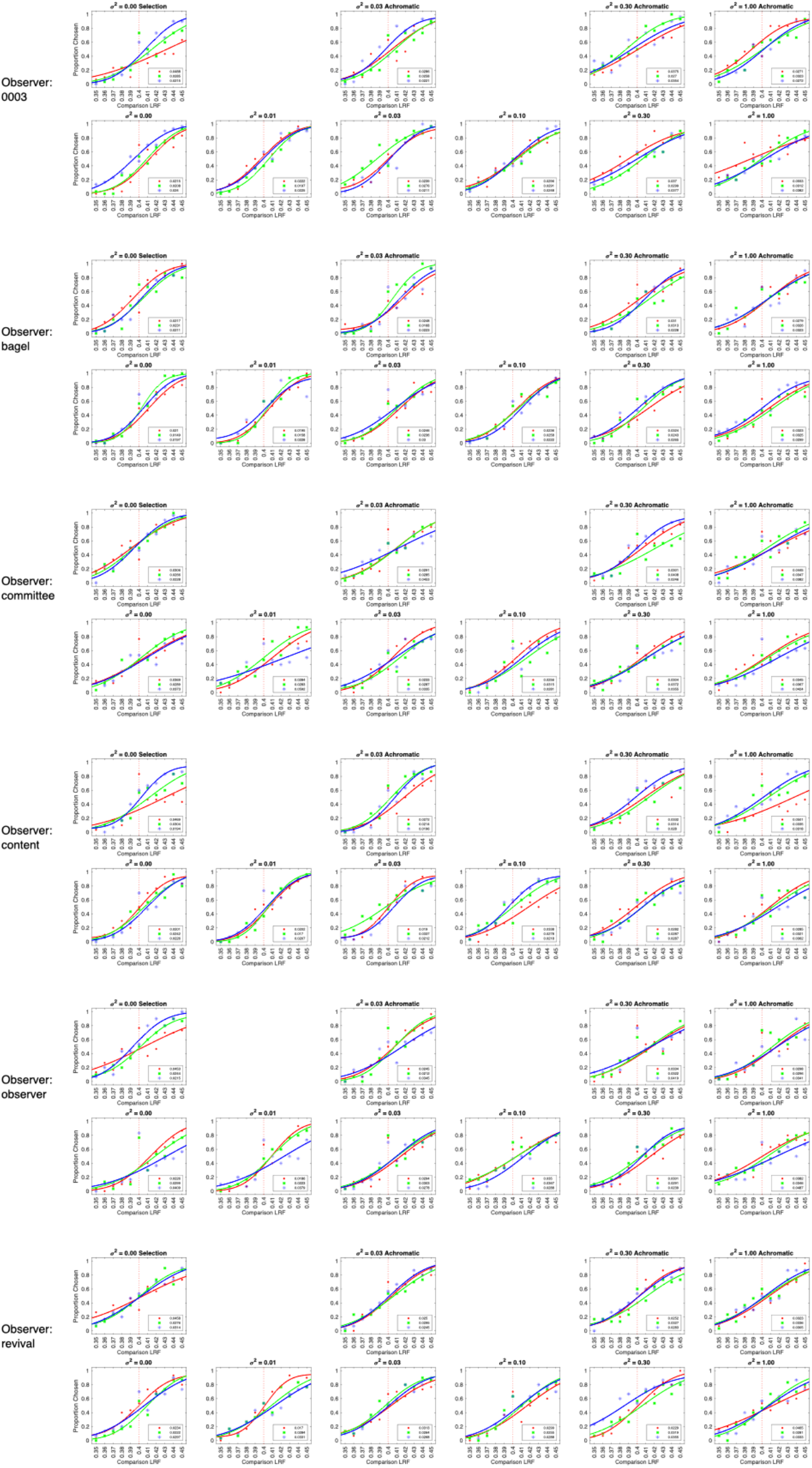
Psychometric functions for all observers for background variation experiment. Same as Figure 6, for all observers retained in the background variation experiment.

**Figure S3:**
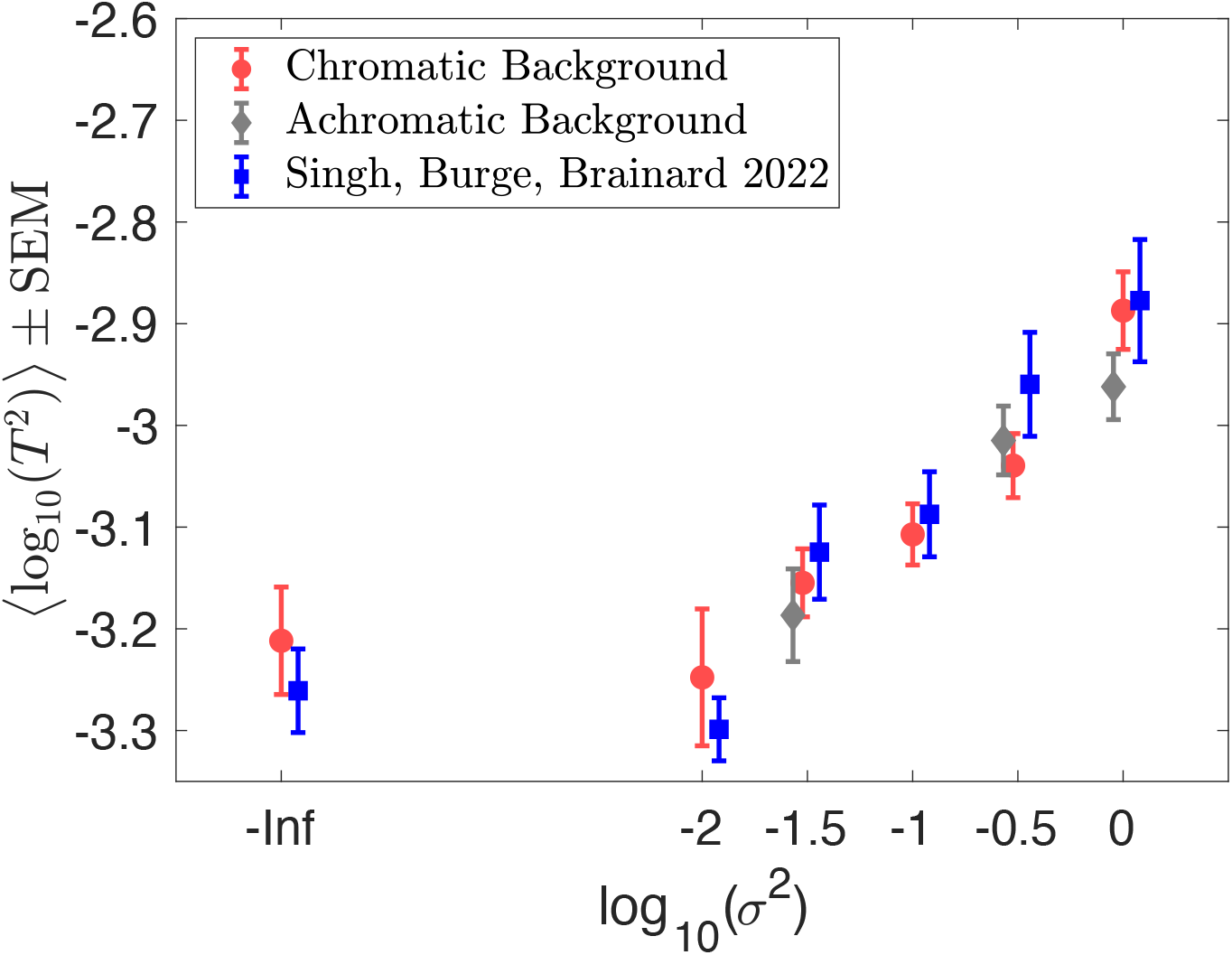
Comparison with Singh, Burge, Brainard 2022. Lightness discrimination thresholds for background variation condition measured in preregistered Experiment 6 and previously reported data from Singh, Burge, Brainard 2022. Preregistered Experiment 6 had both chromatic and achromatic conditions, while the previous experiment (Singh, Burge, Brainard 2022) only had chromatic conditions. Singh, Burge, Brainard 2022 made three threshold measurements for each condition for 4 naïve observers.

**Figure S4:**
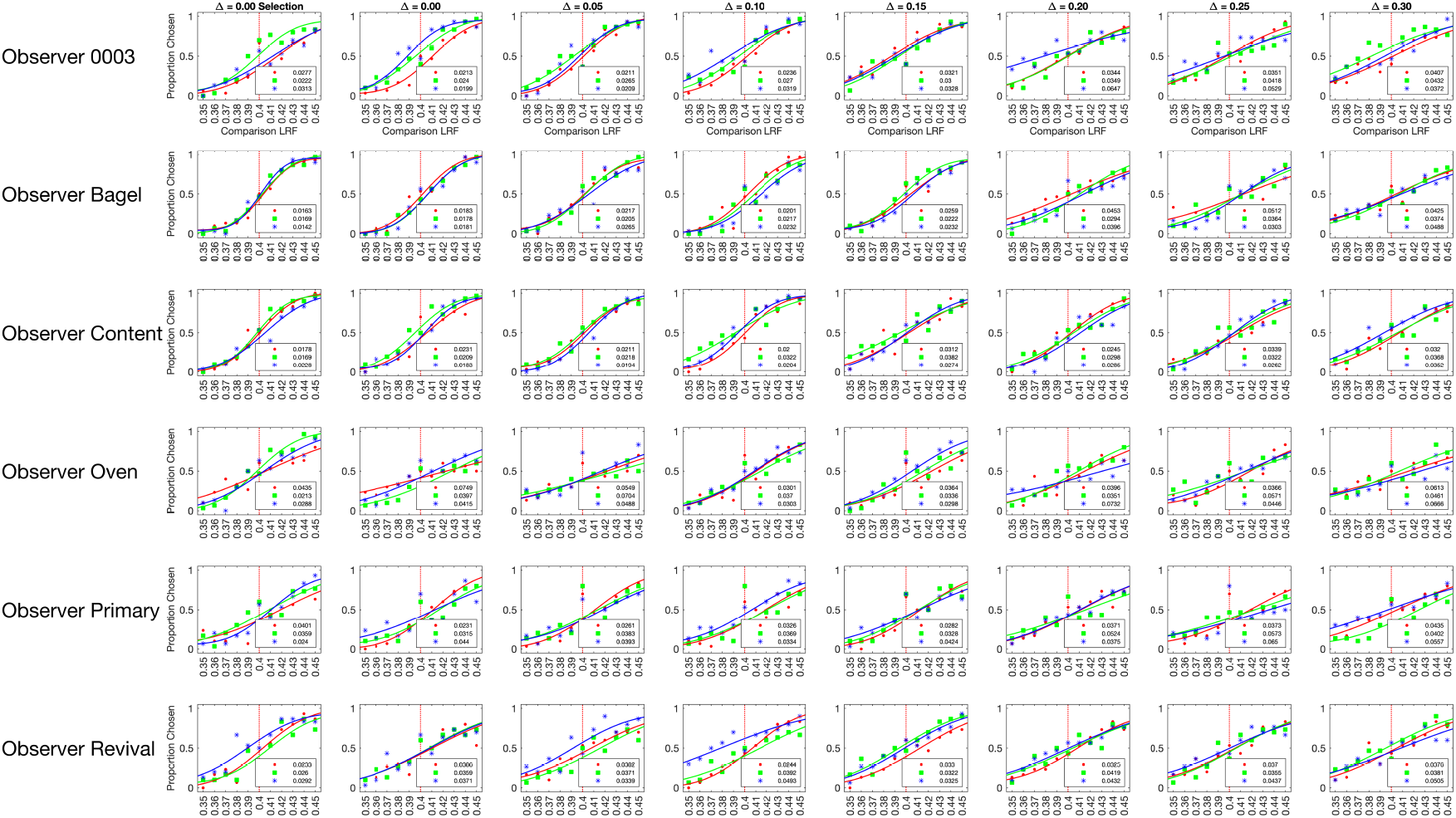
Psychometric functions for all observers for light intensity variation experiment. Same as Figure 8, for all observers retained in the light intensity variation experiment.

**Figure S5:**
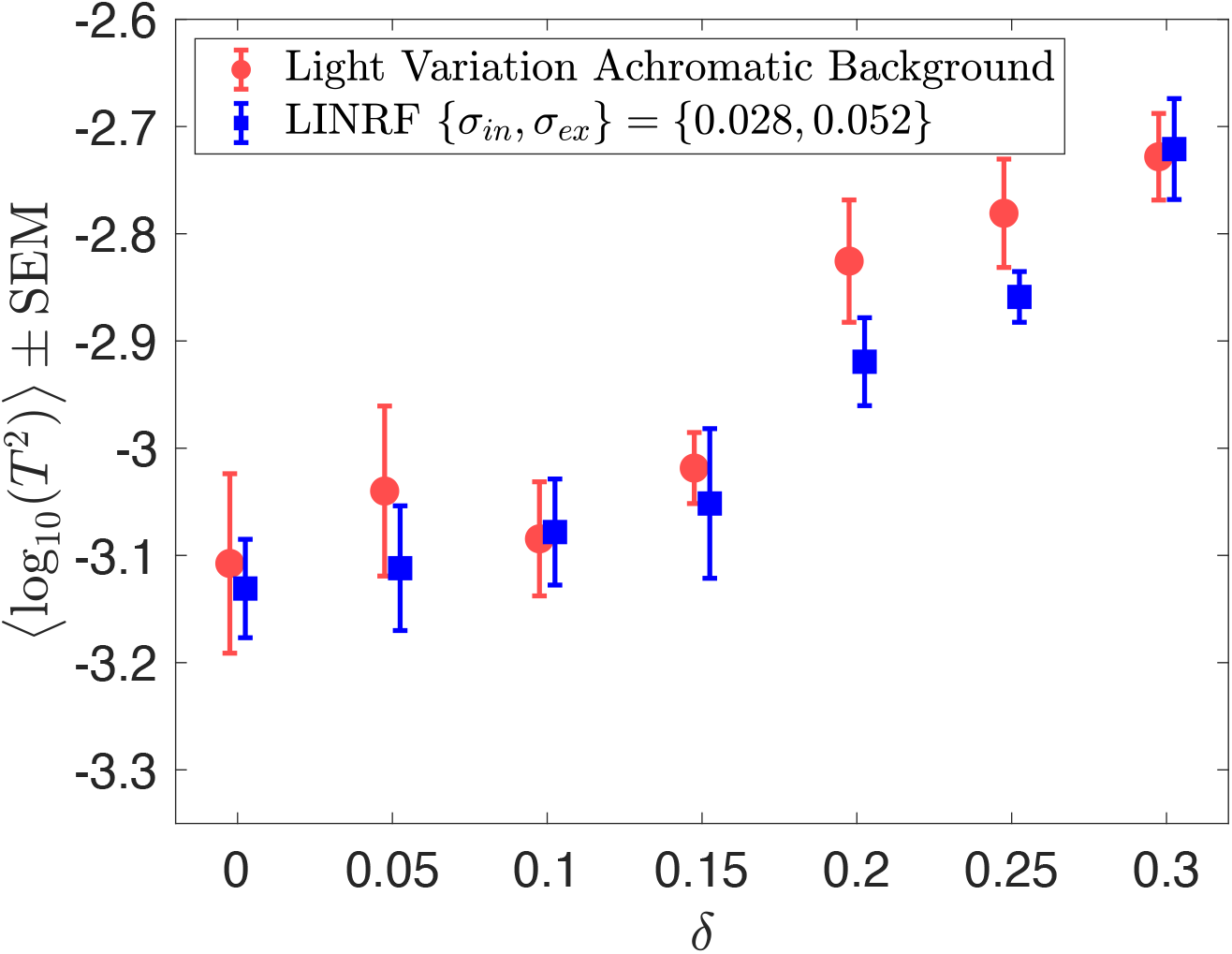
Same as Figure 9, for all six observers retained in the light intensity variation experiment. The parameters for the LINRF model are the same as in Figure 9.

**Figure S6:**
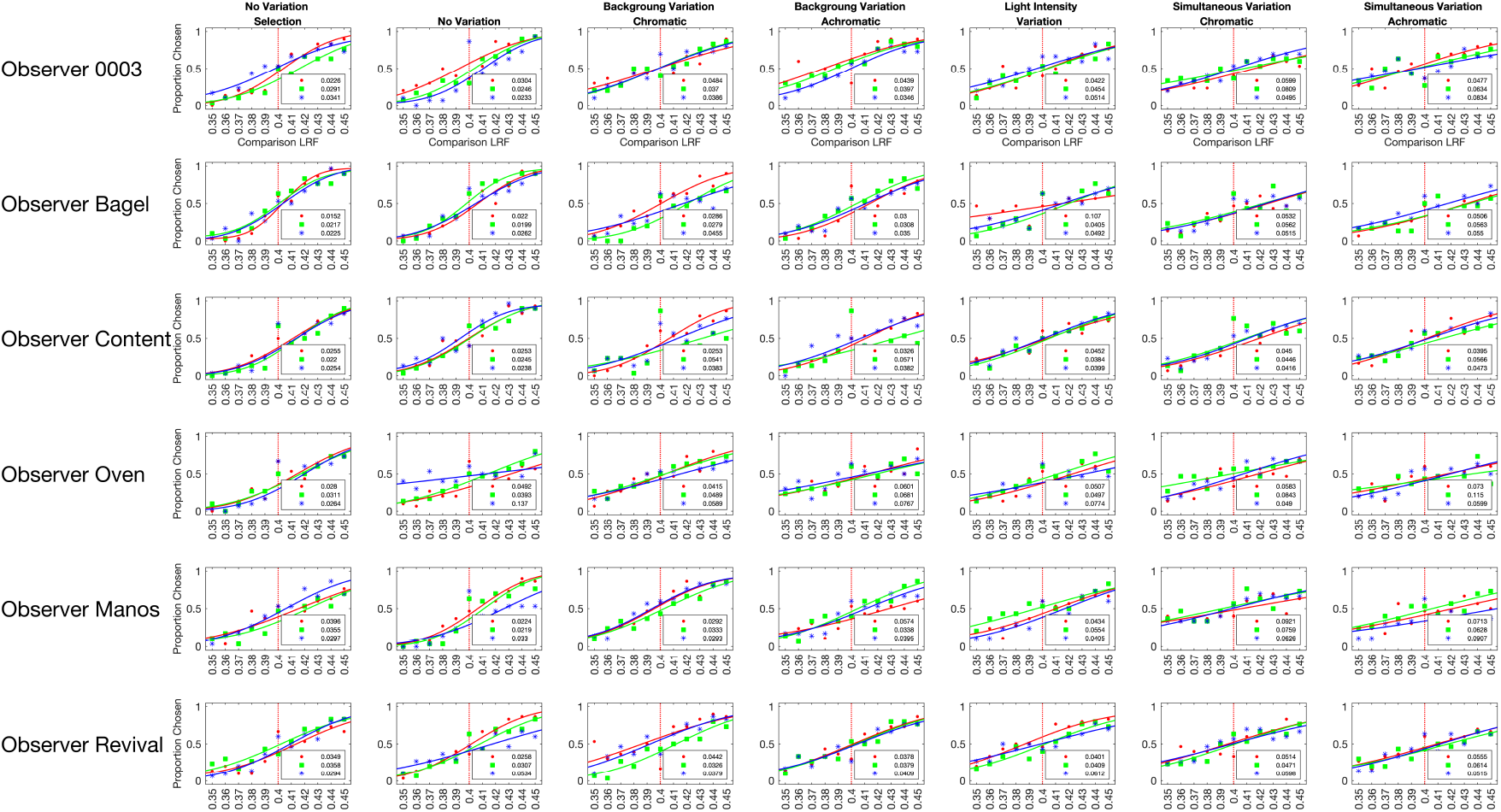
Psychometric functions for all observers for simultaneous variation experiment. Same as Figure 10, for all observers retained in the simultaneous variation experiment.

**Figure S7:**
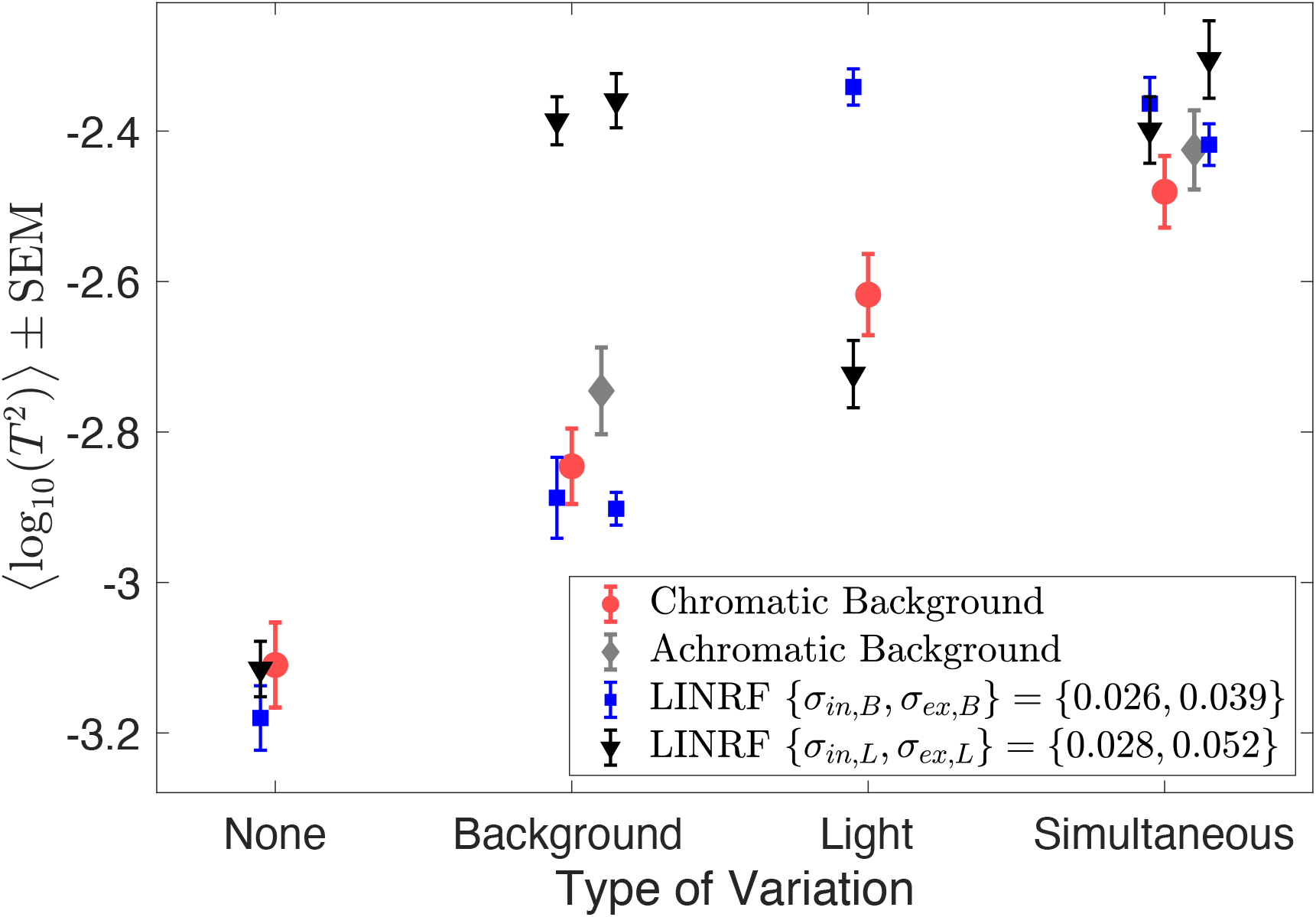
Same as Figure 11, but the thresholds of the linear receptive field (LINRF) model were estimated using the same set of parameters for all six conditions studied in Experiment 8. Blue square markers show log squared thresholds estimated using the parameters of the background variation condition (Experiment 6, Figure 7). Black triangular markers show log-squared thresholds estimated using the parameters of the light intensity variation condition (Experiment 7, Figure 9). The black and blue error bars show +/-1 standard deviation estimated over 10 independent simulations. The parameters of the background variation condition (Experiment 6, blue squares) predict the thresholds of the no variation condition, the background variation condition, and the simultaneous variation condition quite well, but fail to predict the threshold of the light source intensity variation condition. Similarly, the parameters of the light source intensity variation condition (Experiment 7, black triangles) predict the thresholds of the no variation condition, the light source intensity variation condition, and the simultaneous variation condition quite well, but fail to predict the threshold of the background variation condition. This could possibly be because the observers in the three experiments were different. Future work would aim at studying these conditions using the same set of observers.

## Footnotes

1 The preregistration documents relevant to this work are those for Experiments 6, 7 and 8. The site also contains preregistrations for previously reported (Experiment 1, 2 and 3; Singh, Burge, & Brainard, 2022) and unreported (Experiment 4 and 5) work.

